# Intermolecular β-sheet formation guides the interaction between ubiquitin-like modifier FAT10 and adapter protein NUB1L

**DOI:** 10.1101/2025.07.31.667942

**Authors:** Charlotte Weiss, Sarah Overall, Nicola Catone, Alexander B. Barnes, Annette Aichem, Guinevere Mathies

**Affiliations:** Department of Chemistry, University of Konstanz, 78464 Konstanz, Germany; Institute of Molecular Physical Science, ETH Zürich, 8093 Zürich, Switzerland; Institute of Cell Biology and Immunology Thurgau, 8280 Kreuzlingen, Switzerland

**Author notes:** **Corresponding Author** Guinevere Mathies – Department of Chemistry, University of Konstanz, Konstanz 78464, Germany.

**Keywords:** ubiquitin-like modifiers, proteasomal degradation, β-strand capture, fuzzy complexes, magic-angle spinning NMR spectroscopy

## Abstract

Under inflammatory conditions, the ubiquitin-like modifier FAT10 targets proteins for rapid and irreversible degradation by the 26S proteasome. FAT10 is degraded along with its substrates and in this process, the loose folding of FAT10 and adapter protein NUB1L have long been suspected to play crucial roles. We report here the investigation of the N-domain of FAT10 and its interaction with NUB1L by magic-angle spinning (MAS) NMR spectroscopy. A stretch of residues that is intrinsically disordered when the N-domain of FAT10 is in its ubiquitin-like β-grasp fold, becomes part of a regularly structured loop and an intermolecular β-sheet upon binding to NUB1L. The rest of the N-domain is now disordered, with exception of a series of anchor residues and the N-terminus. We propose that, in preparation of degradation by the proteasome, NUB1L stabilizes N-FAT10 in an unfolded state, acting as a holdase. The ability of FAT10 to interact in folded as well as unfolded form is essential for its role in inflammation-linked proteostasis.

## INTRODUCTION

The degradation of proteins into peptides by the 26S proteasome plays a central role in proteostasis and the regulation of cellular pathways, including the immune response.^1,2^ Proteolysis takes place inside the barrel-shaped 20S core of the proteasome, to which access is controlled by the 19S regulatory particle.^3,4^ The small protein ubiquitin serves as the common signal for protein degradation.^5^ Ubiquitin becomes covalently attached to a substrate via an enzyme cascade involving a ubiquitin-activating enzyme (E1), a ubiquitin-conjugating enzyme (E2), and a ubiquitin ligase (E3).^6^ For degradation to proceed, a substrate must have a polyubiquitin chain attached and, in addition, a disordered initiation region^7^. If a substrate is tightly folded, the segregase VCP (valosin containing protein or p97) initiates substrate processing by unfolding a ubiquitin^8^, which is otherwise left intact and reused in the cell.

Ubiquitin-independent pathways of proteasomal degradation also exist. These may be specific for one protein or more generic, such as the midnolin pathway,^9^ which targets transcription factors by β-strand capture. FAT10 (human leukocyte antigen F adjacent transcript 10) is the only other ubiquitin-like modifier (besides ubiquitin itself) that targets substrates directly for degradation by the 26S proteasome.^10,11^ FAT10 consists of two ubiquitin-like domains,^12^ usually referred to as the N- and the C-domain, and it is mainly present in cells and organs of the immune system.^13–15^ In other cell types and tissues, expression is upregulated by the proinflammatory cytokines tumor necrosis factor and interferon-γ.^16,17^ FAT10ylation requires, just like ubiquitylation, a conjugation cascade involving three enzymes: E1 activating enzyme UBA6^18,19^, E2 conjugating enzyme USE1^20^ (several others were recently identified)^21^, and E3 ligase Parkin^22^. Unlike ubiquitin, FAT10 is degraded along with its substrates and, in cells, it has a short half-life of only 1 h.^10^ Interestingly, activation of the proteasome by FAT10 requires a partner, NEDD8-ultimate buster 1 or its long isoform, NUB1(L).^23–25^ FAT10 thus appears to be a tailored tool for a prompt but time-limited response to harmful factors during inflammatory processes.^11^

FAT10 interacts covalently and non-covalently with a diverse set of hundreds of proteins,^26,27^ suggesting a plethora of roles in the immune system. Involvement has been documented in peptide presentation^28^ and autophagy^26^. FAT10 is upregulated in at least a dozen types of cancer, where it is further enhanced by the inflammatory microenvironment of tumors, and has a positive impact on metastasis.^14,29^ The underlying pro-malignant mechanism likely involves binding of FAT10 to the spindle checkpoint protein MAD2 (mitotic arrest deficient 2)^16^; disruption of this binding has been shown to inhibit tumor progression.^30^ At the atomic level, little is known about the binding of FAT10 to its partners and no interaction motifs have been identified. Addressing this knowledge gap is pertinent to the development of therapeutics targeting FAT10 dysregulation in cancer and disorders of the immune response.

FAT10 shows poor solubility and has a tendency to aggregate, even *in vivo*.^31,32^ Indeed FAT10 melts at a relatively low temperature of 41 °C (compared to 83 °C for ubiquitin)^33^ and recent steered molecular dynamics simulations showed low resistance to mechanical unfolding.^34^ The loose folding of FAT10 has long been suspected to be relevant for its biological function, because degradation of FAT10 and FAT10ylated proteins is VCP-independent.^33^ Over the years, it has also interfered with structural investigations. Theng *et al.* determined, by liquid-state NMR, a structure of the N-domain, but could only do so after deletion of seven N-terminal residues.^30^ Later, Aichem *et al.* succeeded in determining a structure of a stabilized, Cys-free mutant of FAT10 by X-ray crystallography (for the N-domain) and liquid-state NMR (for the C-domain).^33^ Both domains are in the β-grasp fold of ubiquitin^35^, but their surfaces differ from ubiquitin and each other. The Cys-free mutant was found to be well conjugated, but, confirming suspicions, was degraded at a much slower pace.

Here, we report the investigation of the Cys-free N-domain of FAT10 (N-FAT10-C0) and its interaction with NUB1L by magic-angle spinning (MAS) NMR spectroscopy. Fast sample spinning around an axis that is at the magic angle of 54.7° with respect to the direction of the magnetic field, makes it possible to acquire high-resolution NMR spectra of biomolecules in the solid state. In this way, inter-atomic distances as well as local chemical environments and dynamics can be probed without restrictions arising from large molecular size or the disorderly arrangement of structural units. This makes MAS NMR well-suited for the atomic-level exploration of interactions between biomolecules, even if one or more binding partners are loosely folded. Recent examples include structural characterization of a branching stimulant that forms condensates upon binding to microtubules, but is disordered in solution^36^ and the observation of a paired helical filament fold of the intrinsically disordered protein tau when bound to lipid membranes^37^.

To start, we sequentially assign the resonances of isolated N-FAT10-C0 to its 83 amino acid residues, based on a series of two- and three-dimensional, ^13^C-detected MAS NMR experiments. The assignment is nearly complete; only the flexible N- and C-terminal tails and residues I68-I74(L76) are absent. From the observed chemical shifts, torsion angles and secondary structure elements are predicted and in agreement with the ubiquitin-like β-grasp fold. Next, we perform the same set of MAS NMR experiments on N-FAT10-C0 in complex with NUB1L. The number of resonances in the spectra is drastically reduced and only residues I68-V81, W17, and the N-terminus of N-FAT10-C0 are sequentially assigned. For an additional eleven amino acids, the type is evident but spectra provide no information about positions in the sequence. Analysis of chemical shifts reveals that residues T73-V81 of N-FAT10-C0 constitute a β-strand and thereby provides experimental evidence for the AlphaFold-Multimer^38^ prediction that the N-domain of FAT10 and NUB1L interact via an intermolecular, anti-parallel β-sheet. With double-REDOR (Rotational-Echo Double-Resonance)^39^ experiments, we confirm that the observed residues of N-FAT10-C0 are at the interface with NUB1L. The disappearance of the remaining residues of N-FAT10-C0 indicates disorder. Hence, we propose that NUB1(L) acts as a holdase for an unfolded N-domain of FAT10, preparing it for rapid degradation by the proteasome.

## RESULTS

### Valuation of Samples

Under the influence of MAS, soluble biomolecules are sedimented, i.e., they become immobilized in a dense phase on the walls of the MAS rotor.^40^ To ensure a high filling factor, this process is usually initiated outside of the spectrometer in an ultracentrifuge, using a dedicated packing tool.^41^ Because of its small size^42^ (9.4 kDa, without isotopic labelling), it is not a priori clear that this sample preparation method is effective for the N-domain of FAT10. We therefore investigated isolated N-FAT10-C0 by MAS NMR in two forms. First, we prepared microcrystals of U-^13^C,^15^N-N-FAT10-C0 using sitting-drop vapor diffusion, with a crystallization solution of ammonium sulfate at pH 2.5. The ^1^H-^13^C and ^1^H-^15^N cross-polarization spectra of microcrystalline N-FAT10-C0 are shown in Figures 1 and S2 (bright green). Conformational homogeneity of N-FAT10-C0 in the microcrystals^43,44^ leads to good resolution, with ^13^C linewidths of 1 ppm or less. Some lines are asymmetric, for example, of ^13^C_δ1_ of Ile at 9 ppm. This is presumably due to the distinct conformations of multiple molecules (A, B, C chains) in the unit cell.

**Figure 1.**
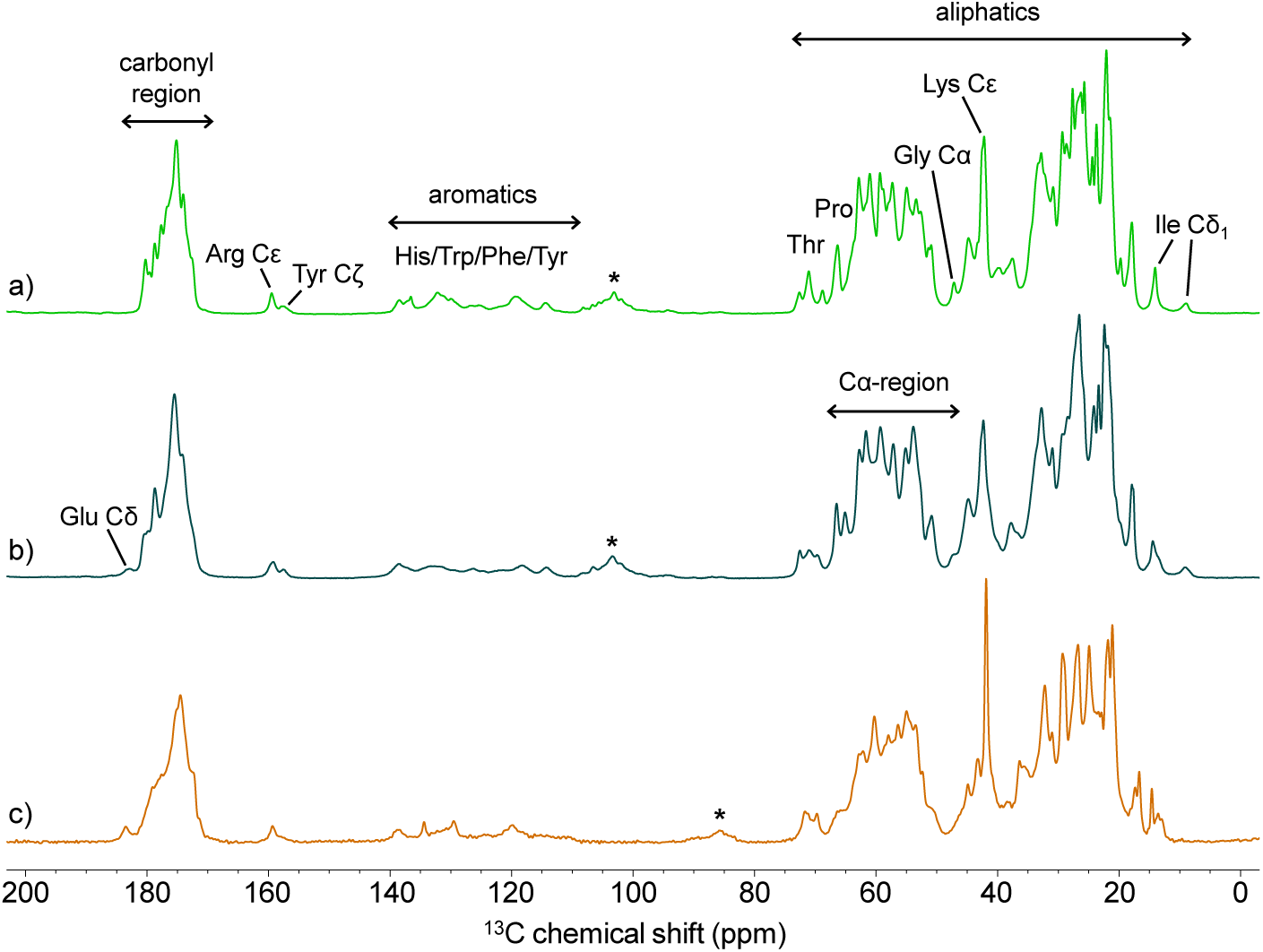
^1^H-^13^C cross-polarization spectra provide fingerprints and an impression of sample integrity. (a) Microcrystalline N-FAT10-C0, (b) lyophilized/rehydrated N-FAT10-C0, and (c) N-FAT10-C0 in complex with NUB1L. Spectra were recorded at 18.8 T with a spinning frequency of 14.5 kHz (a,b) and at 20.0 T with a spinning frequency of 19.0 kHz (c). The sample temperature was 4 °C. The stars indicate spinning sidebands.

Second, we prepared a non-crystalline reference sample at near-physiological pH. To this end, U-^13^C,^15^N-N-FAT10-C0 was lyophilized from buffer solution (pH 7.5), ground, and manually packed into a MAS rotor. To rehydrate, deionized water was added in steps of 1 μL until spectral resolution no longer improved. The resulting ^1^H-^13^C and ^1^H-^15^N cross-polarization spectra of lyophilized N-FAT10-C0 are shown in Figures 1 and S2 (turquoise) and are very similar to those of microcrystalline N-FAT10-C0. Linewidths have increased (in spite of rehydration), but the C_α_-region remains resolved and prominent resonances from aromatic sidechains and ^13^C_δ1_ of Ile remain in place. The altered appearance of the carbonyl region is due to the change in pH: at pH 7.5, Asp and Glu are deprotonated and their ^13^C_γ_ and ^13^C_δ_ resonances have shifted downfield^45,46^.

Prior to investigation by MAS NMR, we verified that not only the wild-type^47^, but also the stabilized, Cys-free variant of the N-domain of FAT10 used in this study forms a complex with NUB1L. Figure S1 shows the positive result, from size-exclusion chromatography. In the MAS rotor, the complex was formed using co-sedimentation^48,49^ by ultracentrifugation from a buffer solution (pH 7.5) containing U-^13^C,^15^N-N-FAT10-C0 and natural abundance NUB1L in a 1:1 molar ratio. The spectra of N-FAT10-C0 co-sedimented with NUB1L (orange in Figures 1 and S2) show well-defined resonances, particularly from aromatic and aliphatic sidechains. At the same time, the C_α_-region is not as nicely resolved as in the spectra of isolated N-FAT10-C0. Most strikingly, compared to these spectra, features have drastically changed.

### Resonance Assignments and Secondary Structure of Isolated N-FAT10-C0

Figure 2a shows the aliphatic region of the ^13^C-^13^C correlation spectrum of microcrystalline N-FAT10-C0, recorded with the dipolar assisted rotational resonance (DARR) pulse sequence^50^ (the full ^13^C-^13^C spectrum is shown in Figure S3a). With a mixing time of 10 ms, the spectrum is dominated by one- and two-bond transfers, which enables convenient identification of individual amino acid residue types based on sidechain patterns. To find the positions of the amino acids in the sequence, we recorded a ^15^N-^13^C-^13^C correlation spectrum with the *z*-filtered transferred echo double-resonance (ZF TEDOR) DARR pulse sequence^51,52^. This sequence transfers magnetization from ^15^N nuclei to ^13^C nuclei going forward (NCACX) and backward (NCOCX) along the backbone simultaneously. A TEDOR mixing time of ∼1 ms emphasizes one-bond (N-C_α_ and N-C’) magnetization transfers. Thereafter, a DARR mixing time of 40-50 ms allows transfer from C_α_ or C’ nuclei to other ^13^C nuclei roughly within a sidechain. A representative strip plot of the ZF TEDOR DARR spectrum of microcrystalline N-FAT10-C0 (Figure 2b) illustrates the efficient magnetization transfer and the large number of resolved cross peaks. A ^15^N-^13^C correlation spectrum, recorded with the ZF TEDOR sequence alone, offers an overview of the backbone connections and supports the identification of residues with a ^15^N nucleus in the sidechain (Figure S3b).

**Figure 2.**
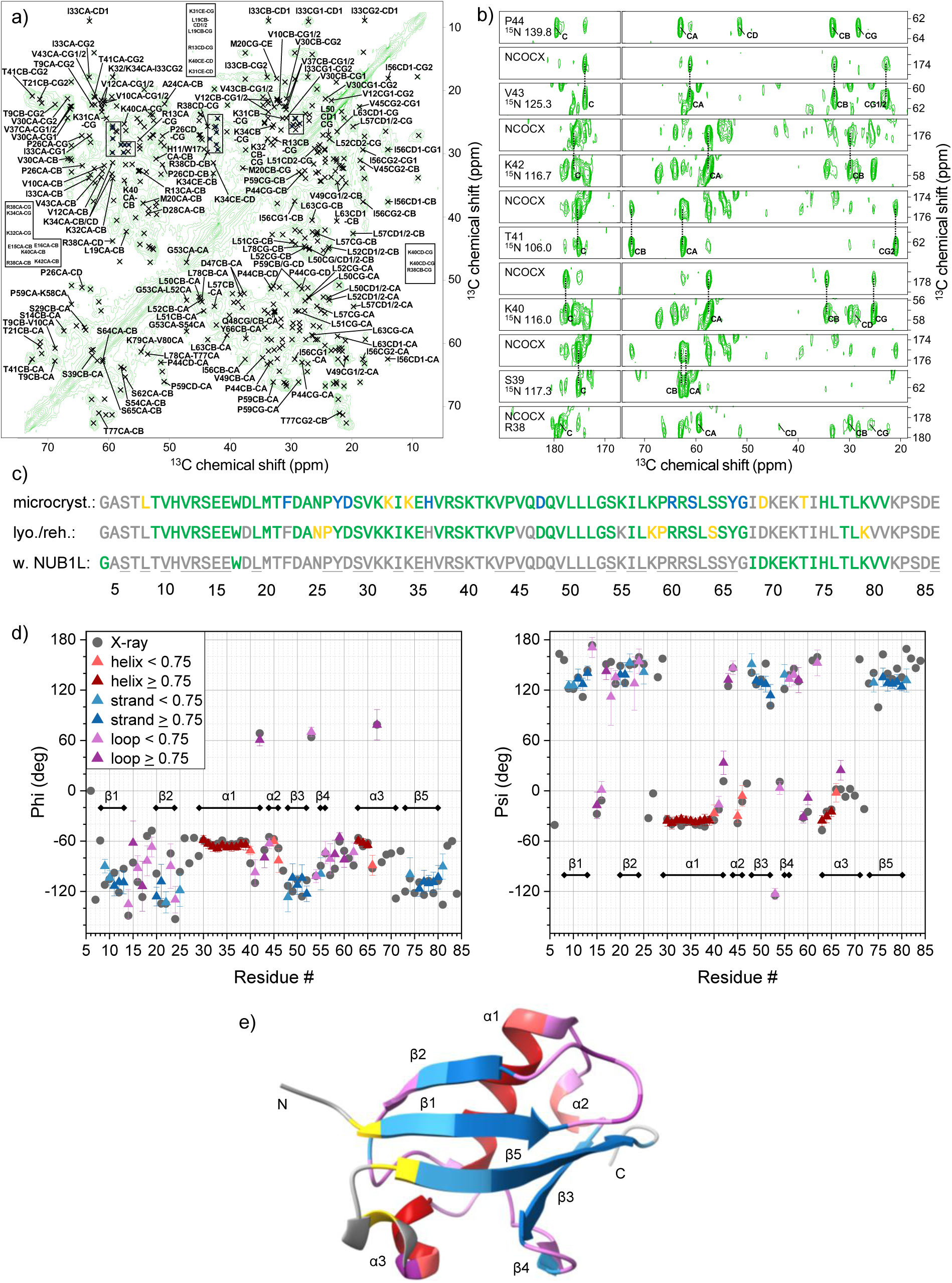
MAS NMR spectroscopy yields reliable resonance assignments and secondary structure predictions for N-FAT10-C0. (a) Aliphatic region of the ^13^C-^13^C DARR spectrum of microcrystalline N-FAT10-C0. Sequentially assigned cross peaks are marked and labeled (residues 4-43 above the diagonal, 44-86 below the diagonal). (b) Representative strip plot, from the ^15^N-^13^C-^13^C ZF TEDOR DARR spectrum of microcrystalline N-FAT10-C0, showing the backbone walk from R38 to P44. (c) Overview of sequential assignments of microcrystalline N-FAT10-C0, lyophilized/rehydrated N-FAT10-C0, and N-FAT10-C0 in complex with NUB1L. Green = unambiguous, yellow = ambiguous, blue = multiple conformations, grey = unobserved. Of the grey, underlined residues of N-FAT10-C0 in complex with NUB1L, we observe 1×E, 1×I, 1×L, 1×P, 1×R, 2×S, 3×V, and 1×Y, but sequential assignment was not possible. (d) Backbone torsion angle (triangles) and secondary structure predictions based on MAS NMR of microcrystalline N-FAT10-C0. Error bars correspond to the standard deviation of the Φ and Ψ angles of the best matches in the TALOS-N database. For comparison, torsion angles (black dots) and secondary structure elements of the X-ray structure of N-FAT10-C0 (PDB-ID 6GF1, chain B) are also plotted. Predictions with a confidence ≥ 0.75 are highly reliable. See the Supporting Information for a more detailed description of the TALOS-N results. (e) Projection of predicted secondary structure per residue on the X-ray structure of N-FAT10-C0.

With these three spectra in hand, we were able to sequentially assign 69 of the 83 residues of N-FAT10-C0. Chemical shifts are listed in Table S3 and Figure 2c provides an overview. Unobserved residues (grey) are restricted to the flexible N- and C-terminal tails (G4-T7 and K82-E86) and the stretch I68-I74. Figure 2d shows predictions for the backbone torsion angles Φ and Ψ per residue based on the observed chemical shifts. The colours indicate the predicted secondary structure (red for helix, blue for strand, and purple for loop). The torsion angles of the X-ray structure of N-FAT10-C0^33^ are also plotted (black dots). Agreement with predicted angles is very good; the root of the mean square deviation is 14° for Φ and 12° for Ψ. In Figure 2e, predicted secondary structure elements are projected onto the X-ray structure. Agreement is again very good, demonstrating that, with the set of MAS NMR experiments we chose^53,54^, reliable resonance assignments and secondary structure predictions are obtained for the N-domain of FAT10.

The aliphatic region of the ^13^C-^13^C correlation spectrum of lyophilized/rehydrated N-FAT10-C0 is shown in Figure 3a (the full ^13^C-^13^C and the ^15^N-^13^C correlation spectra are shown in Figure S4). As expected from the one-dimensional spectra, resolution is not as good as for microcrystalline N-FAT10-C0, but, in the ^13^C-^13^C spectrum, sidechain patterns are still recognizable. In Figure 3b, a representative strip plot of the ^15^N-^13^C-^13^C correlation of lyophilized/rehydrated N-FAT10-C0 is shown. Transfer of magnetization along the backbone is less efficient with the ZF TEDOR DARR sequence than for the microcrystalline sample. To address this issue, a supplementary NCOCX spectrum was recorded, with a narrow spectral width and an increased number of scans, using SPECIFIC-CP^55^ and combined 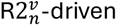 (CORD)^56^ ^13^C-^13^C mixing.

**Figure 3.**
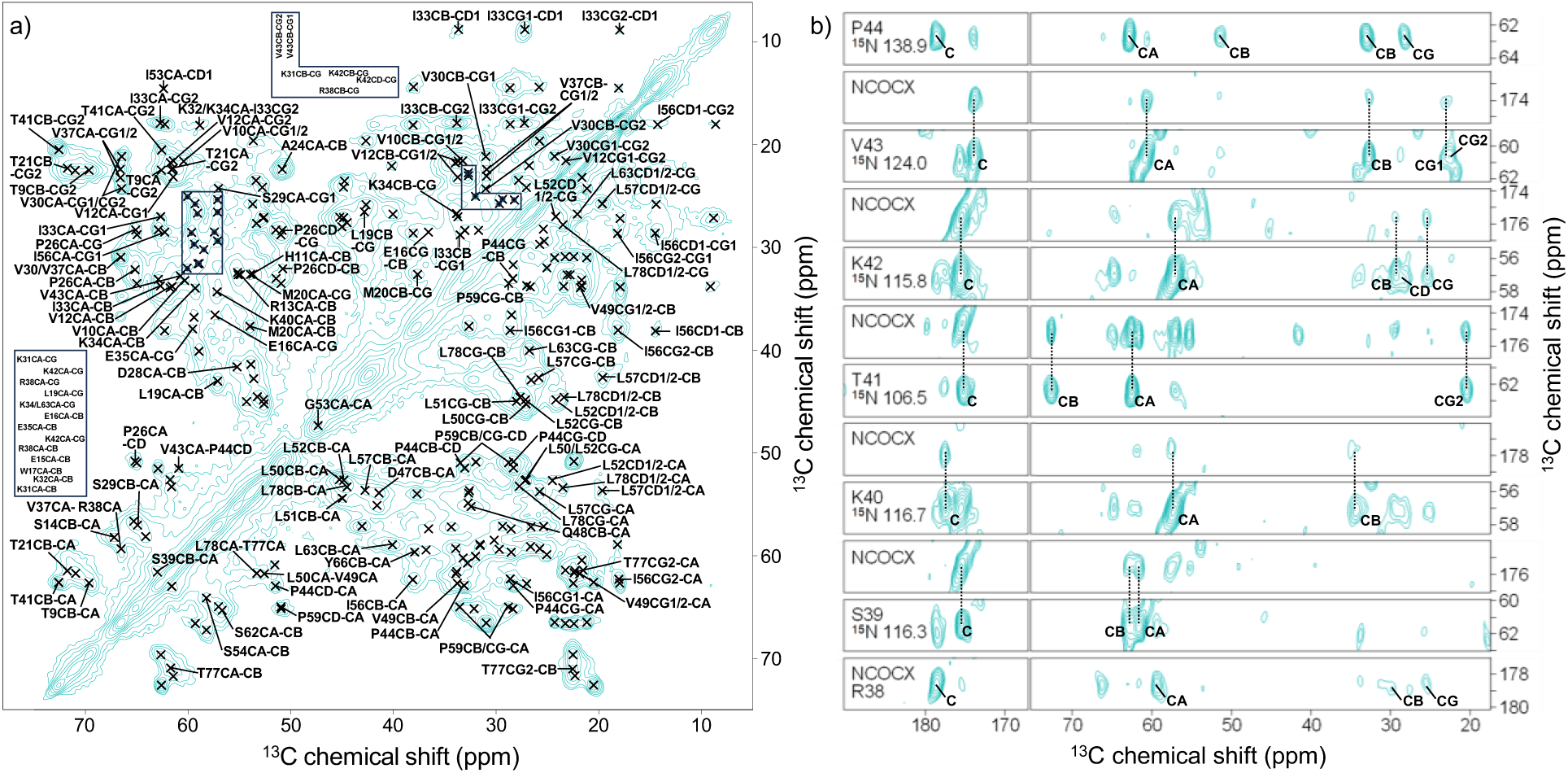
MAS NMR spectra of lyophilized/rehydrated N-FAT10-C0. (a) Aliphatic region of the ^13^C-^13^C DARR spectrum. Sequentially assigned cross peaks are marked and labeled (residues 4-43 above the diagonal, 44-86 below the diagonal). (b) Representative strip plot, from the ^15^N-^13^C-^13^C ZF TEDOR DARR/SPECIFIC CP CORD spectra of lyophilized/rehydrated N-FAT10-C0, showing the backbone walk from R38 to P44.

The resonance assignment of lyophilized/rehydrated N-FAT10-C0 was performed *de novo*, but the known chemical shifts of microcrystalline N-FAT10-C0 provided essential guidance. For example, the sidechain patterns of I33 and I56 are very similar in both samples. In total, we sequentially assigned 56 of the 83 residues (Figure 2c and Table S4). Similarly to microcrystalline N-FAT10-C0, residues in the flexible tails (G4-L8 and V80-E86) are not observed nor is the stretch I68-L76. In addition, six residues scattered along the sequence are not visible. Comparison of the chemical shifts of the microcrystalline and lyophilized/rehydrated samples of N-FAT10-C0 shows small differences (1-2 ppm), which tend to be larger, up to 5 ppm, if a neighboring residue is not observed in the lyophilized/rehydrated sample. Possibly multiple conformations of (neighboring) bulky sidechains create local disorder in the backbone. Vice versa, C_α_-N cross peaks of W17 and G67 were not observed for the microcrystalline sample, but are visible in the spectra of the lyophilized/rehydrated sample (in Figure S4b, at 59.1,119.0 ppm and 45.9,107.1 ppm), again suggesting local changes in structure. Predicted torsion angles and secondary structures are, however, very similar for both samples of isolated N-FAT10-C0, indicating that the ubiquitin-like β-grasp fold is intact (Figure S5).

### Interaction of N-FAT10-C0 with NUB1L

Figures 4a and b show the aliphatic (and aromatic) regions of the ^13^C-^13^C and ^15^N-^13^C correlation spectra of N-FAT10-C0 co-sedimented with NUB1L (full spectra in Figure S6). Well-resolved cross peaks are observed, but, compared to the spectra of isolated N-FAT10-C0 (Figures 2a, 3a, S3, and S4), their number is drastically reduced. The same is true for the ^15^N-^13^C-^13^C correlation spectra (ZF TEDOR DARR and additional NCOCX) of the co-sediment. A possible reason for the disappearance of signals from the spectra could be large-amplitude motion. The NMR pulse sequences we used are designed for the investigation of solids, which means that they start with a ^1^H-^13^C cross-polarization step and rely on steady dipolar couplings between the nuclei for magnetization transfer. Modulation of dipolar couplings interferes with this transfer, possibly rendering parts of N-FAT10-C0 invisible in the spectra. To investigate if this is happening here, we recorded a series of one-dimensional, directly-excited ^13^C spectra at temperatures of 4, -2 and -10 °C (Figure S7). Comparison to spectra recorded with ^1^H-^13^C cross-polarization at the same temperatures (Figure S8) reveals modest changes in relative intensities, but no new signals. The only effect of lowering the temperature is a slight decrease in the resolution. We surmise that not large-amplitude motion, but disorder (which may still be dynamic, see the Discussion) has broadened signals from a large part of N-FAT10-C0 beyond detection.

**Figure 4.**
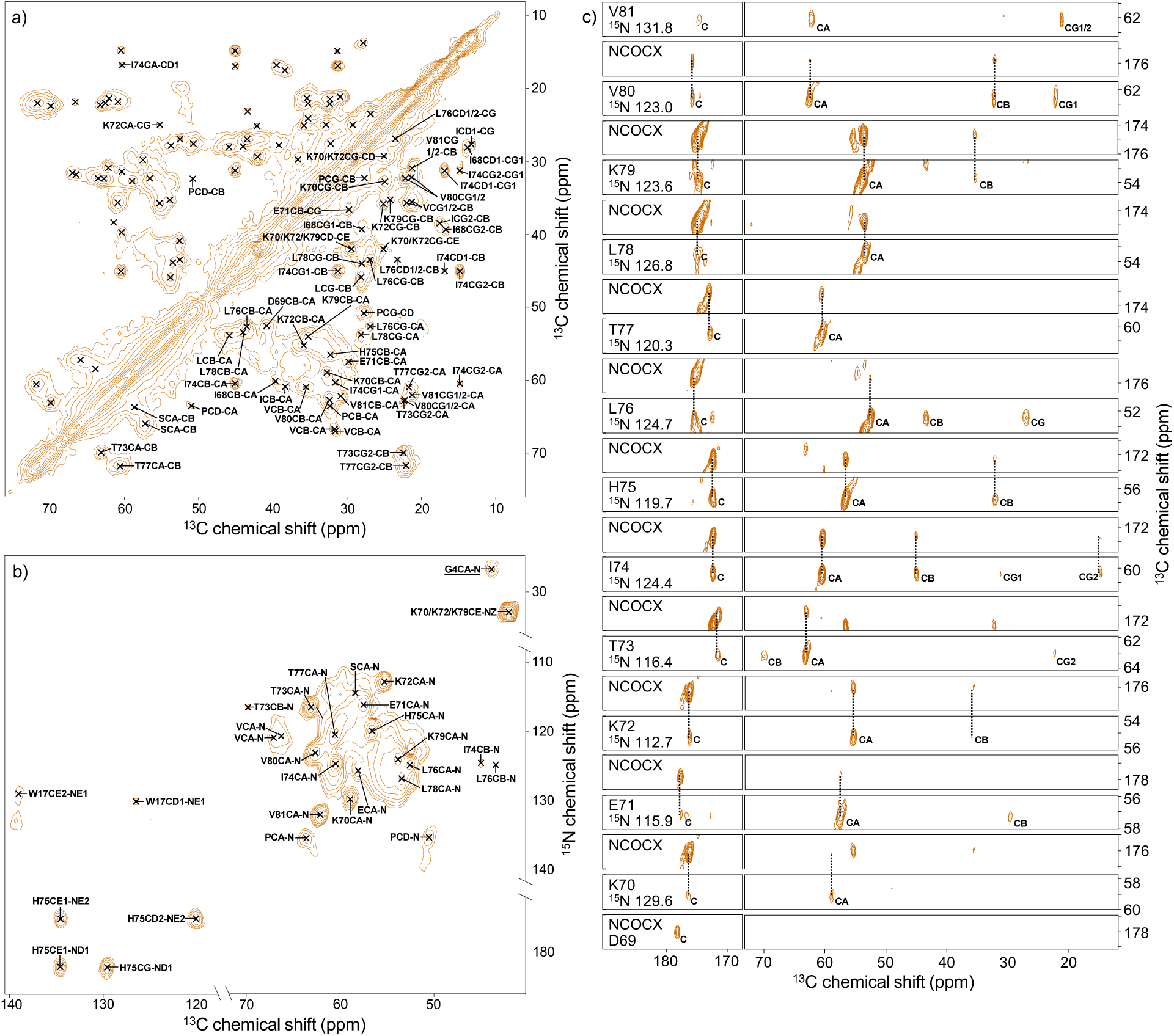
MAS NMR spectra of N-FAT10-C0 co-sedimented with NUB1L. (a) Aliphatic region of the ^13^C-^13^C DARR spectrum. Sequentially assigned cross peaks are marked and labeled. (b) Aliphatic and aromatic regions of the ^15^N-^13^C ZF TEDOR spectrum. (c) Strip plot showing the backbone walk from D69 to V81. Strips are from the ^15^N-^13^C-^13^C ZF TEDOR DARR spectrum, except for the NCOCX strips of D69-T73, which are from the SPECIFIC CP CORD spectrum. The N and C’ nuclei of I68 are not observed, but proximity to D69 was confirmed by Cγ2-Cα and Cγ2-Cβ cross peaks in DARR spectra with long mixing times.

Analysis of the signals that *are* observed for the co-sediment of N-FAT10-C0 and NUB1L, leads to a confident sequential assignment of residues I68-V81 (Figure 2c). The strip plot that demonstrates the magnetization transfers along the backbone, is shown in Figure 4c. For another twelve amino acids (1×E, 1×I, 1×L, 1×P, 1×R, 2×S, 3×V, 1×W, 1×Y), sidechain patterns are evident, but magnetization transfer to neighboring residues is not detected. Hence, the position of these residues in the amino acid sequence remains unknown, except for W17, which is the only Trp in the sequence. Intriguingly, a pronounced C_α_-N cross peak is observed in the ^15^N-^13^C correlation spectrum (Figure 4b) at 43.8, 26.8 ppm. Given the unusual ^15^N chemical shift, this cross peak must arise from the N-terminus of N-FAT10-C0, G4. A corresponding C’-C_α_ cross peak of this Gly is visible at 169.6, 43.8 ppm in the ^13^C-^13^C correlation spectrum (Figure S6a). A table of all observed chemical shift values is provided in the Supporting Information (Table S5). Torsion angle and secondary structure predictions based on the chemical shifts of the sequentially assigned residues are plotted in Figure S9. Residues I68-K72 form a loop, while residues T73-V81 form a β-strand.

To further explore the interaction between N-FAT10-C0 and NUB1L, we used structure prediction by AlphaFold-Multimer. The best-ranked structure is shown in Figure 5a (confidence scores are provided in Figure S10). N-FAT10-C0 is in an open, partially unfolded state and NUB1L is wrapped around an intermolecular, anti-parallel β-sheet that consists of residues L76-V81 of N-FAT10-C0 and Y212-N217 of NUB1L. This predicted structure suggests a plausible interpretation of the MAS NMR experimental data. In preparation of rapid degradation by the proteasome, interaction with NUB1L stabilizes N-FAT10-C0 in an unfolded, mostly disordered state. Only the residues visible in the spectra have retained a well-defined, stable structure. If they could be sequentially assigned, they are highlighted in orange in Figure 5a.

**Figure 5.**
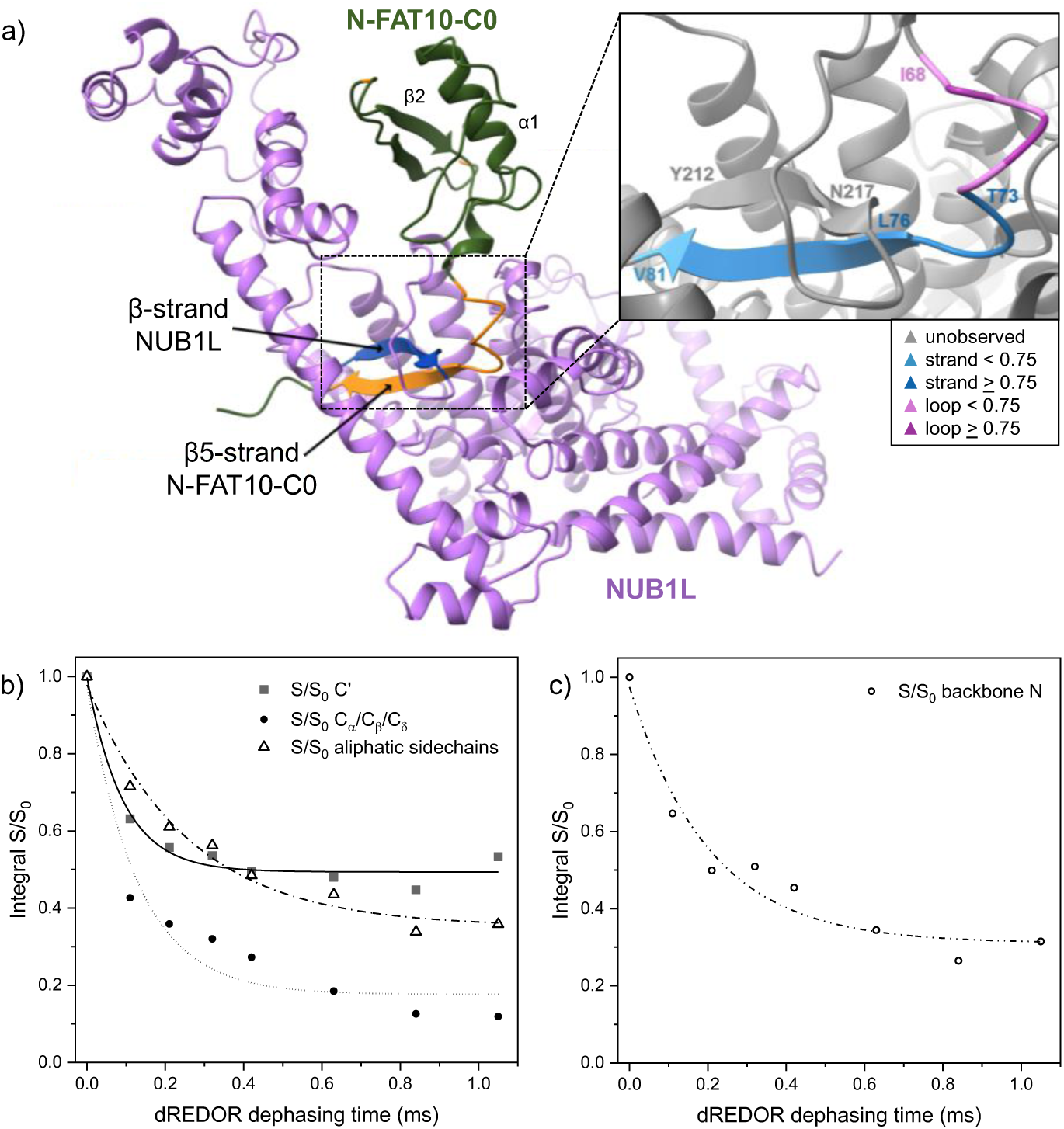
Prediction and experimental observation of the interface between N-FAT10-C0 and NUB1L. (a) AlphaFold-Multimer structure prediction for the complex of N-FAT10-C0 and NUB1L. The residues of N-FAT10-C0 that are observed by MAS NMR are highlighted in orange. The inset shows the empirical secondary structure predictions for residues I68-V81 projected on the AlphaFold-Multimer structure. See Figure S9 for the backbone torsion angles and details. MAS NMR confirms the existence of the intermolecular β-sheet and shows that it is in reality three residues longer. Integrity of the other secondary structure elements in N-FAT10-C0 is negated by MAS NMR. (b,c) Double-REDOR dephasing dynamics for different types of ^13^C and backbone ^15^N nuclei of N-FAT10-C0 co-sedimented with NUB1L. Exponential decay functions fitted to the experimental data points highlight that ^1^H magnetization has been replenished after double-REDOR dephasing. See Figures S11-S13 for one-dimensional spectra and details of the analysis.

The inset in Figure 5a shows the empirical secondary structure predictions, based on observed chemical shifts, projected on the AlphaFold-Multimer structure. MAS NMR thus provides experimental evidence for the formation of the intermolecular β-sheet. In reality, it is even three residues longer: on the side of N-FAT10-C0, residues T73-V81 participate not just L76-V81. The residues of N-FAT10-C0 preceding the β-strand form a loop until I68, again in agreement with the AlphaFold-Multimer structure. MAS NMR, however, does not confirm the structural integrity of the β2-strand and α1-helix of N-FAT10-C0, which AlphaFold-Multimer predicts with high confidence (Figures 5a and S10). Instead, twelve residues scattered along the N-FAT10-C0 chain plus the Gly N-terminus are observed. We suspect that these residuesform anchor points where N-FAT10-C0 interacts non-covalently with NUB1L in between disordered stretches.

To explore the interactions between U-^13^C,^15^N-N-FAT10-C0 and natural-abundance NUB1L, we used the double-REDOR (Rotational-Echo Double-Resonance) filter^39^. In this MAS NMR experiment, following pre-saturation of ^13^C (or ^15^N) nuclei and a 90°-pulse on protons, the magnetization of protons directly bonded to ^13^C and ^15^N nuclei is dephased by the simultaneous application of ^1^H-^13^C and ^1^H-^15^N REDOR pulses. Proton magnetization remaining on natural-abundance NUB1L is subsequently transferred via cross-polarization to ^13^C (or ^15^N) nuclei on U-^13^C,^15^N-N-FAT10-C0. In a one- or two-dimensional spectrum that is finally recorded, only ^13^C (or ^15^N) nuclei at the interface with NUB1L will be visible.

We recorded one-dimensional ^1^H-^13^C and ^1^H-^15^N cross-polarization spectra with the double-REDOR filter for both the reference sample of lyophilized/rehydrated N-FAT10-C0 and for N-FAT10-C0 co-sedimented with NUB1L. Spectra were obtained with the double-REDOR pulses switched off and on, for a series of dephasing times, see Figures S12 and S13. Decay of signals is evident due to the natural loss of ^1^H spin coherence and due to double-REDOR dephasing. For each of the spectra, we quantified signal intensities (S_0_ without and S with the double-REDOR pulses) of the C’-region, the C_α_-region (including C_β_ of Ser and Thr and C_δ_ of Pro), the sidechain carbons, and backbone nitrogens. As expected, and in agreement with the results of Polenova *et al.* for unbound proteins,^39^ the signals of lyophilized/rehydrated N-FAT10-C0 are fully dephased after approximately 0.8 ms (Figure S11). For N-FAT10-C0 co-sedimented with NUB1L, however, dephasing remains incomplete, indicating that ^1^H magnetization is replenished and transferred from nearby NUB1L (Figures 5b,c). For the C’-region, the sidechain carbons, and the backbone nitrogens, the ratio S/S_0_ is 0.4-0.5 after more than 1 ms of dephasing, again in agreement with the results obtained by Polenova *et al.* for a test protein bound to its partner^39^. For the C_α_-region, the final ratio S/S_0_ is clearly less, about 0.1. One possible cause may be a short T_1ρ_ in combination with a relatively large distance between the C_α_-nuclei of N-FAT10-C0 and the undephased protons of NUB1L, e.g., in an anti-parallel β-sheet inter-strand hydrogen bonds are formed between carbonyls and amines resulting in ^1^H-C_α_ distances of ∼4 Å *vs.* ^1^H-C’ distances of ∼3 Å.

To trace the incomplete dephasing back to individual residues, we recorded a two-dimensional DARR spectrum, preceded by double-REDOR dephasing. This yielded a ^13^C-^13^C correlation spectrum of N-FAT10-C0 co-sedimented with NUB1L with a much cleaned up diagonal in which signals from virtually all previously observed residues of N-FAT10-C0 can be identified, see Figure S14. In addition, cross peaks are present from all sequentially assigned residues, except W17, D69, and H75, and from non-sequentially assigned residues Leu, Ser, and Val. Taken together, the experiments with the double-REDOR filter support our conjecture that the residues of N-FAT10-C0 observed by MAS NMR (Figure 4) owe their stable structures to close contact with NUB1L.

## DISCUSSION

The E1 enzyme UBA6 is unusual in the sense that it not only activates FAT10, but also ubiquitin.^18,19^ Co-crystallization of UBA6 with Cys-free FAT10 has shown that the binding of the C-domain of FAT10 is analogous to the binding of ubiquitin to the, ubiquitin-only, E1 enzyme UBA1.^57^ The N-domain of FAT10 interacts with the three-helix bundle of UBA6, with the β-grasp fold intact. The conjugation cascade thus requires FAT10 in properly folded form. The interaction of N-FAT10 with UBA6 bears similarity with the interaction of another ubiquitin-like modifier NEDD8 (neural precursor cell-expressed developmentally down-regulated protein 8) with NUB1L: both interactions require a ubiquitin-associated (UBA) domain, are polar, and rely on N51 of NEDD8 or the analogous residue K58 in N-FAT10. Based on this information, we had naively expected N-FAT10-C0 to remain folded upon binding to NUB1L. To assess the feasibility of a MAS NMR study, we first prepared a microcrystalline sample of isolated N-FAT10-C0. Aiming to gain insight into the binding mode and interface via chemical shift perturbations, we also required the lyophilized/rehydrated reference sample of N-FAT10-C0 at physiological pH – fortunately its preparation and characterization by MAS NMR were successful. Resonance assignment of N-FAT10-C0 co-sedimented with NUB1L, however, constituted a plot twist by revealing a drastic change in the fold of the N-domain of FAT10.

Last year, Arkinson *et al.* reported the investigation of the conformation and solvent accessibility of wild-type FAT10 upon binding with NUB1 (the shorter splice variant of NUB1L) using hydrogen-deuterium exchange detected by mass spectrometry.^58^ In the absence of NUB1, several peptides from both the N- and the C-domain show a bimodal distribution, indicating coexisting folded and unfolded states. In the presence of NUB1, residues throughout the N-domain were exposed, except for the last β5-strand. The authors concluded that NUB1 traps the unfolded N-domain of FAT10. A coexistence of folded and unfolded states may well explain the troubles encountered in the investigation of the N-domain of FAT10 by liquid-state NMR. Theng *et al.* were only successful after deletion of the first seven N-terminal residues and even then residues P59-T73 could not be detected by ^1^H-^15^N heteronuclear single-quantum correlation (HSQC); in the structure they obtained, the α1-helix is somewhat displaced (PDB-ID 2MBE).^30^

Arkinson *et al.* found their conclusion supported by structure prediction with AlphaFold-Multimer and by site-directed mutagenesis: FAT10 mutants H75A, H75D and H75K compromised NUB1 binding, H75D and H75K also abolished degradation.^58^ Cao *et al.* recently went a step further showing that phosphorylation of T77 interferes with NUB1L binding.^59^ Arkinson *et al.* detected no complex formation between full-length Cys-free FAT10 and NUB1, while in our hands complex formation between N-FAT10-C0 and NUB1L is evident (Figure S1). Possibly the absence of the C-domain (in our case) makes it easier for the N-domain to become inserted into the clasp of NUB1L. Finally, Arkinson *et al.* showed by cryo-electron microscopy that FAT10 binding induces an open conformation of NUB1, allowing its ubiquitin-like (UBL) domain to interact with the RPN1 subunit^60^ of the regulatory particle for direct FAT10 delivery. In spite of directed efforts, however, it was not possible to resolve the interaction of NUB1 with FAT10.

In biomolecular MAS NMR, the observation of an amino acid residue is indicative of a well-defined, stable structure. In the case of N-FAT10-C0 co-sedimented with NUB1L, this applies to residues I68-V81, of which I68-K72 form a loop and T73-V81 form a β-strand, the N-terminus (G4), W17, and eleven more residues that could not be sequentially assigned (Figure 2c). The other, unobserved residues of N-FAT10-C0 evade detection due to disorder. This disorder is likely not static, but dynamic, meaning that these residues sample a large, heterogeneous conformational space, probably on a slow to intermediate timescale (milli- to microseconds). In recent years, integrated approaches of MAS NMR, liquid-state NMR, relaxation dispersion, and molecular dynamics simulations have cast light on such dynamic disorder in proteins and enzymes. The dynamic nature of intrinsically disordered or structurally frustrated regions are critical for function, enabling, for example, interactions with other biomolecules and catalytic activity^61,62^.

Based on the selective survival of residues in the MAS NMR spectra^63^, the AlphaFold-Multimer prediction (see below), and the results previously obtained by Arkinson *et al.*, we propose that the N-domain of FAT10 and NUB1(L) form a fuzzy complex^64^. More specifically, NUB1(L) is a predominantly folded chaperone for the predominantly unfolded client N-FAT10(-C0). Unlike for isolated N-FAT10-C0, the signals of N-FAT10-C0 co-sedimented with NUB1L are retained after double-REDOR dephasing. This confirms the existence of an interface between the two proteins, consisting of the intermolecular β-sheet, the preceding loop region, isolated anchor residues scattered along the amino acid chain, and the N-terminus. ^13^C and ^15^N nuclei in the sidechains of the anchor residues are more consistently observed than their backbone counterparts (Table S5), suggesting that these residues engage in electrostatic and hydrophobic interactions with NUB1L predominantly via their sidechains. Electrostatic interactions probably also play a role in the binding of the N-terminus. The N-terminus is known to enter the proteasome first^10,33,58^ and, hence, control over its position and orientation is likely critical for the presentation of unfolded FAT10 to the proteasome. In addition, the stabilization of N-FAT10 in its folded form upon deletion of the N-terminal tail (Theng *et al.*)^30^ hints at a possible role of the binding of the N-terminus in the unfolding of N-FAT10 by NUB1L. Figure 6 summarizes what we know about the fuzzy complex of N-FAT10 and NUB1(L).

**Figure 6.**
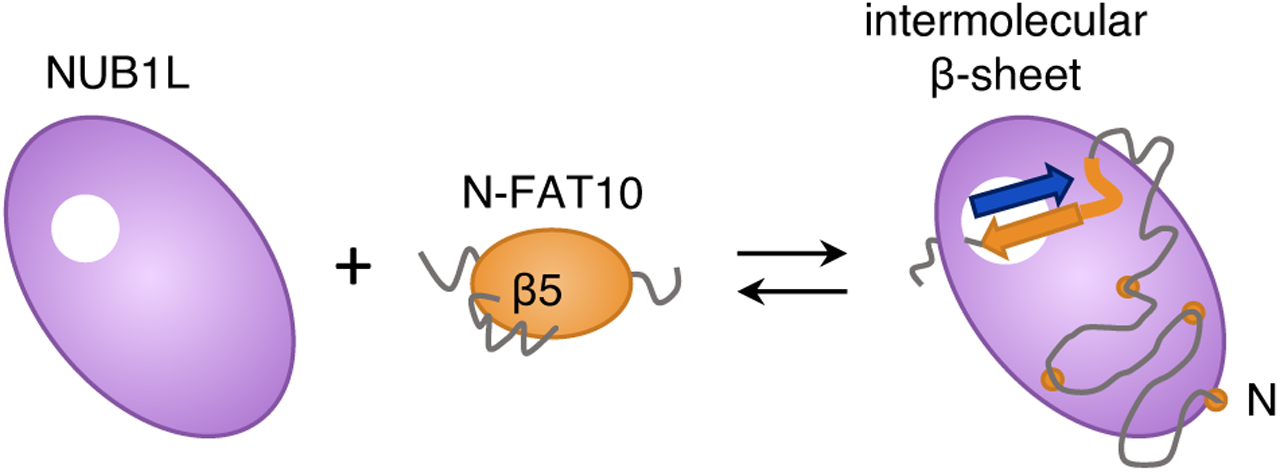
N-FAT10(-C0) and NUB1L form a fuzzy complex. When the N-domain of FAT10 is in the ubiquitin-like β-grasp fold, residues I68-I74(L76) are disordered and primed for β-strand capture. Upon interaction with NUB1L, residues I68-K72 form an ordered loop and T73-V81 become part of an intermolecular, anti-parallel β-sheet. The rest of N-FAT10 is now disordered, with exception of a series of anchor residues and the N-terminus (indicated by the orange circles). NUB1L acts as a holdase, stabilizing a largely unfolded N-FAT10.

Structure-based sequence alignment of FAT10 with ubiquitin reveals two extra residues in the N-domain preceding the β5-strand (K70, E71), extending the loop. Molecular dynamics simulations of the N-domain in its β-grasp fold have previously indicated loose folding in this region.^33^ This is in agreement with our experimental observations by MAS NMR: residues I68-I74(L76) were not or barely observed (Figure 2c), indicating disorder. Multiple sequence alignment of mammalian FAT10 proteins reveals a highly conserved surface comprising residues T73, H75, L76, T77, L78, and V80. The disordered region of folded N-FAT10 is thus primed for formation of the intermolecular β-sheet with NUB1L, see Figure 6. The proceeding is reminiscent of the capture of an unstructured β-strand degron by two Catch subdomains in the midnolin pathway of ubiquitin-independent protein degradation.^9^ In the case of N-FAT10, β-strand capture appears to constitute an interaction motif. Titrations in combination with liquid-state NMR have implicated the β1- and β5-strands of N-FAT10 in the binding with MAD2.^30^ MAD2, in turn, is known to undergo β-sheet augmentation with reshuffling upon complex formation with CDC20 (cell-division-control protein 20) or MAD1.^65^ This unusual interaction motif, however, brings the risk of aggregation. In β2-microglobulin, for example, the pathogenic D76N mutation destabilizes protective edge β-strands, which exposes aggregation-prone regions and enables amyloid formation.^66^

AlphaFold-Multimer predicts with high confidence that the β2-strand and α1-helix of N-FAT10-C0 remain partially intact (Figures 5a, S10), but this is not reflected in the MAS NMR spectra. This likely relates to the known tendency of AlphaFold to predict, for intrinsically disordered proteins or regions, the structures of conditionally folded states, i.e., stable structures that form under specific conditions such as the interaction with a binding partner or following post-translational modification.^67^ For example, AlphaFold’s predicted structure of isolated 4E-BP2 simultaneously contains the four-β-strand structure that forms upon multisite phosphorylation^68^ and two helices that resemble those observed in crystal structures of 4E-BP2 and 4E-BP1 bound to translation initiation factor 4E.^69,70^ The prediction of the intact β2-strand and α1-helix is thus likely a remnant of the conditional folding of isolated N-FAT10-C0. There is additional disagreement between AlphaFold-Multimer and the MAS NMR data regarding the length of the intermolecular β-sheet. AlphaFold-Multimer predicts residues L76-V81 of N-FAT10-C0 to participate whereas TALOS-N classifies T73-V81 as β-strand. This is not just due to the rendering of the secondary structure: the backbone torsion angles of I74 are clearly distinct (TALOS-N: Φ = −133.3°, Ψ = 155.3°; AlphaFold-Multimer/VADAR: Φ = −117.1°, Ψ = −24.2°, see Figure S9). To accommodate the longer β-sheet, NUB1L would have to undergo a conformational change. Finally, AlphaFold-Multimer does not predict the interactions of NUB1L with the scattered anchor residues of N-FAT10-C0, nor the binding of the N-terminus. All this emphasizes that experimental data remain essential in the investigation of the structures and functions of loosely folded (or intrinsically disordered) proteins. MAS NMR experiments that could, in combination with suitable isotopic labelling, provide further information about the fuzzy complex of N-FAT10 and NUB1L include frequency-selective rotational-echo double-resonance (REDOR)^71^ to measure specific ^13^C-^15^N distances, possibly across the interface, ^1^H detection enabled by ultrafast MAS^72^ to explore the hydrogen bonding network of the anti-parallel β-sheet, and relaxation dispersion^73^ to probe microsecond timescale conformational sampling.

## CONCLUSION

Investigation with MAS NMR of the ubiquitin-like modifier FAT10 and its interaction with adapter protein NUB1L has captured a built-in shapeshifting of the N-domain of FAT10 at the atomic level. To become covalently attached to its substrates, both the N- and the C-domain of FAT10 must be in the ubiquitin-like β-grasp fold. In this situation, a stretch of residues leading up to the β5-strand of the N-domain experiences structural frustration, which makes them a target for β-strand capture by NUB1L. Formation of the intermolecular β-sheet with NUB1L forces this stretch of residues into a regular structure, which is incompatible with the β-grasp fold. NUB1L becomes a holdase for the N-domain of FAT10, stabilizing it in a mostly unfolded state for rapid and VCP-independent degradation of FAT10 and its substrates. The ability of FAT10 to interact with binding partners both in folded and in unfolded form is key to its mediation of proteasomal degradation.

## MATERIALS and METHODS

Below we give short accounts. Details can be found in the Supporting Information.

### Expression and Purification of U-^13^C,^15^N-N-FAT10-C0

Cys-free N-domain of human FAT10 (amino acids 5-86; C7T, C9T) was expressed as a His_6_-GST-fusion protein in *E. coli* BL21-CodonPlus(DE3)-RIPL competent cells (Agilent Technologies). An additional glycine residue at the N-terminus remains from the TEV protease cleavage site. For uniform ^13^C and ^15^N labelling, cells were grown in M9 minimal medium supplemented with U-^13^C_6_-D-glucose and ^15^N-ammonium chloride. The purification was adapted from a previously published protocol.^33^ Briefly, bacterial cells were lysed and the supernatant was applied to nickel affinity chromatography. After buffer exchange and cleavage of the purification tag, His-tagged TEV protease and byproducts were separated by a second nickel affinity chromatography. Final purification by size-exclusion chromatography yielded up to 7 mg per litre of M9 medium.

### Expression and Purification of NUB1L

Human NUB1L (amino acids 2-615) was expressed as a His_6_-SUMO-fusion protein, also in *E. coli* BL21-CodonPlus(DE3)-RIPL competent cells. To produce natural abundance NUB1L, cells were grown in LB medium. Purification requires the same steps as for U-^13^C,^15^N-N-FAT10-C0. Yield was up to 15 mg per litre of LB medium.

### MAS NMR Spectroscopy

Experiments were performed at 18.8 and 20.0 T (^1^H Larmor frequencies of 800 and 850 MHz) on Bruker Avance NEO/III spectrometers, each equipped with a 3.2 mm E-free HCN Bruker MAS probe. The spinning frequency was 14.5 or 19.0 kHz. The sample temperature was 4 °C, unless noted otherwise. Chemical shifts of ^13^C are referenced to DSS in D_2_O (0.5% by weight), chemical shifts of ^15^N are referenced to liquid ammonia at 25 °C. Protocols for sample packing, an overview of the MAS NMR experiments (Table S1), and all acquisition parameters (Table S2) are provided in the Supporting Information. Spectra were processed in TopSpin or NMRPipe^74^. CcpNmr Analysis^75^ was used for the resonance assignments.

### Torsion Angle and Secondary Structure prediction

Backbone torsion angles Φ and Ψ and secondary structure were predicted empirically based on chemical shifts of assigned N, C’, C_α_, and C_β_ nuclei.^76,77^ For this purpose, we used TALOS-N^78^, which combines a set of trained neural networks with efficient mining of a database of proteins of which both the X-ray structure and the chemical shifts are known.

### AlphaFold Modelling

The structure of isolated N-FAT10-C0 (amino acids 5-86 with C7T and C9T and the additional glycine at the N-terminus) was predicted with AlphaFold^79^ v2.3.2 (reduced database). The maximum template release date was 2014-08-26. Five predictions were obtained, from one seed per model, and ranked according to the predicted local distance difference test (pLDDT) confidence score. Only the best prediction was relaxed. The structure of the complex of N-FAT10-C0 and NUB1L (amino acids 1-615) was predicted with AlphaFold-Multimer (reduced database).^38^ The maximum template release date was 2004-11-28. Twenty-five predictions were obtained, from five seeds per model, and ranked according to a weighted combination of the predicted template modelling (pTM) and interface pTM scores (model confidence = 0.8·ipTM + 0.2·pTM). Again, only the best prediction was relaxed. All AlphaFold modelling was performed on the Scientific Compute Cluster of the University of Konstanz (SCCKN). UCSF ChimeraX^80^ was used for structure visualization. Torsion angles were extracted from the AlphaFold structures using VADAR^81^.

## ASSOCIATED CONTENT

### Supporting Information

Expression and purification protocols for U-^13^C,^15^N-N-FAT10-C0 and NUB1L; preparation of samples for MAS NMR, including size-exclusion chromatography and SDS-PAGE showing complex formation of N-FAT10-C0 and NUB1L; overview of all MAS NMR experiments and acquisition parameters; ^1^H-^15^N cross-polarization spectra; ^13^C-^13^C and ^15^N-^13^C spectra, representative strip plot, table with ^15^N and ^13^C chemical shifts, and details of torsion angle analysis of microcrystalline N-FAT10-C0; representative strip plots, table with ^15^N and ^13^C chemical shifts, and torsion angle and secondary structure predictions of lyophilized/rehydrated N-FAT10-C0; direct excitation ^13^C and ^1^H-^13^C cross-polarization spectra at various temperatures, strip plot, table with ^15^N and ^13^C chemical shifts, and torsion angle and secondary structure predictions of N-FAT10-C0 in complex with NUB1L; AlphaFold-Multimer confidence scores; one-dimensional ^13^C and ^15^N double-REDOR spectra of lyophilized/rehydrated N-FAT10-C0 and analysis; one-dimensional ^13^C and ^15^N double-REDOR spectra and ^13^C-^13^C double-REDOR DARR spectrum of N-FAT10-C0 in complex with NUB1L.

### Data availability

All data generated in this study have been deposited in the <u>Zenodo repository</u>. The NMR assignments for microcrystalline N-FAT10-C0, lyophilized/rehydrated N-FAT10-C0, and N-FAT10-C0 in complex with NUB1L have been deposited to the Biological Magnetic Resonance Data Bank (BMRB) under accession codes 53484 and 53485.

## AUTHOR INFORMATION

### Notes

The authors declare no competing financial interest.

## Acknowledgments

The authors are deeply indebted to Prof. Dr. Marcus Groettrup. Without his enthusiasm and scientific excellence, this work would not have been possible. He is sorely missed. This research was funded by the Deutsche Forschungsgemeinschaft through the Emmy Noether Program (Project No. 321027114), SFB 969 (Project No. 189682160), and SFB 1527 (Project No. 454252029) and by the Swiss National Science Foundation (Projects No. 201070 and 219514). The authors thank Michael Kovermann, Ulrich Haunz, and Anke Friemel for maintenance of the NMR instrumentation, Venkata SubbaRao Redrouthu for support with the measurements, Leon Schöwe and Gunter Schmidtke for support in the biolab, Salima Bahri for practical advice, and Stefan Gerlach for support regarding the Scientific Compute Cluster at the University of Konstanz (SCCKN).

