## Supporting Information for "Intermolecular β-sheet formation guides the interaction between ubiquitin-like modifier FAT10 and adapter protein NUB1L"

### 1. Protein expression and purification

**U-$^{13}$C,$^{15}$N-N-FAT10-C0**

Cysteine-free N-domain of human FAT10 (amino acids 5-86; C7T, C9T) was expressed as a $His_6$-GST-fusion protein in *E. coli* BL21-CodonPlus(DE3)-RIPL competent cells (Agilent Technologies). The pETM-30 plasmid was kindly provided by the Institute of Cell Biology and Immunology Thurgau (Kreuzlingen, Switzerland); a TEV protease cleavage site C-terminal to the purification tag results in an additional glycine residue at the N-terminus after cleavage. For uniform $^{13}$C and $^{15}$N labelling, 3.6 g of U-$^{13}C_6$-D-glucose and 0.5 g of $^{15}$N-ammonium chloride per litre of M9 minimal medium were added as exclusive sources of carbon and nitrogen. Bacteria cells were grown at 37 °C to an $OD_{600}$ of 0.5-0.6, induced with 0.4 mM IPTG at 21 °C overnight, and harvested by centrifugation (RCF 4000 g, 10 min, 8 °C). Harvested cells were lysed in lysis buffer (20 mM TRIS-HCl (pH 8.0), 300 mM NaCl, 10 mM imidazole, 10 % v/v glycerol, 0.1 % v/v Triton X-100, 100 µg/mL lysozyme, 1 mM PMSF and EDTA-free protease inhibitor). After sonication, cell debris was removed by centrifugation (RCF 47000 g, 30 min, 8 °C). The supernatant was filtered and applied to Ni-NTA Agarose beads (Macherey-Nagel) for affinity chromatography. The protein was eluted with elution buffer (20 mM TRIS-HCl (pH 8.0), 300 mM NaCl and 500 mM imidazole) and buffer exchanged to binding buffer (20 mM TRIS-HCl (pH 8.0), 300 mM NaCl and 10 mM imidazole). His-tagged TEV protease was allowed to act overnight (5 µg of protease per 1 mg of recombinant protein). After cleavage, His-tagged TEV protease and cleavage byproducts were separated by a second Ni-affinity chromatography using binding buffer. The volume of the flow-through containing the U-$^{13}$C,$^{15}$N-N-FAT10-C0 was reduced to 5 mL and filtered. Further purification was achieved by size-exclusion chromatography (Cytiva, formerly Amersham Biosciences, ÄKTA pure chromatography system equipped with a HiLoad 16/600 Superdex 75 pg column) yielding up to 7 mg per litre of M9 medium. Before preparation of the samples for MAS NMR, U-$^{13}$C,$^{15}$N-N-FAT10-C0 was in 20 mM HEPES (pH 7.5) and 150 mM NaCl and concentrated to specified values (Amicon Ultra-15 centrifugal filter unit, MWCO 3 kDa).

**NUB1L**

Human NUB1L (amino acids 2-615) was expressed as a $His_6$-SUMO-fusion protein also in *E. coli* BL21-CodonPlus(DE3)-RIPL competent cells. The pSUMO plasmid was kindly provided by the Institute of Cell Biology and Immunology Thurgau (Kreuzlingen, Switzerland). Bacteria cells were grown in LB medium at 37 °C to an $OD_{600}$ of 0.7, induced with 0.4 mM IPTG at 21 °C overnight and harvested by centrifugation (RCF 4000 g, 10 min, 8 °C). The harvested cells were lysed in lysis buffer (20 mM TRIS-HCl (pH 7.5), 150 mM NaCl, 20 mM imidazole, 1 mM TCEP, 10 % v/v glycerol, 0.1 % v/v Triton X-100, 100 µg/mL lysozyme, 1 mM PMSF and EDTA-free protease inhibitor). After sonication, cell debris was removed by centrifugation (RCF 27000 g, 30 min, 8 °C). The supernatant was filtered and applied to Ni-NTA Agarose beads for affinity chromatography. The protein was eluted with elution buffer (20 mM TRIS-HCl (pH 7.5), 150 mM NaCl, 500 mM imidazole and 1 mM TCEP) and buffer exchanged to binding buffer (20 mM TRIS-HCl (pH 7.5), 150 mM NaCl, 20 mM imidazole and 1 mM TCEP). His-tagged Ulp1 protease was added overnight (4 µg of protease per 1 mg of recombinant protein). After cleavage, His-tagged Ulp1 protease and cleavage byproducts were separated by a second Ni-affinity chromatography using binding buffer. The volume of the flow-through containing (natural abundance) NUB1L was reduced to 5 mL and filtered. Further purification was achieved by size-exclusion chromatography yielding up to 15 mg of pure NUB1L per litre of LB medium. To form the complex with N-FAT10-C0, the protein was in 20 mM HEPES (pH 7.5), 150 mM NaCl, and 1 mM TCEP, concentrated to approximately 20 mg/mL (Amicon Ultra-15 centrifugal filter unit, MWCO 30 kDa).

#### 2. Packing of samples for MAS NMR

**Microcrystalline N-FAT10-C0**

Microcrystals of U-$^{13}C$,$^{15}N$-N-FAT10-C0 were grown following an adapted version of the crystallization protocol of Aichem *et al.*[1], with the sitting-drop vapor diffusion method. Equal volumes of protein at a concentration of approximately 20 mg/mL and crystallization solution (3.2 M $(NH_4)_2SO_4$ and 0.1 M citric acid (pH 2.5)) were mixed in drops of 25 µL. Each drop was equilibrated against 0.75 mL of crystallization solution supplemented with additional 150 mM NaCl. After keeping the crystallization trays at 4 °C for four days, microcrystals were harvested. Ultracentrifugation was used to assure dense packing in a 3.2 mm MAS rotor. A home-made packing tool[2] containing the protein suspension and MAS rotor was centrifuged for 2 h at RCF 210000 g and 4 °C in a Beckman Coulter Optima L90K ultracentrifuge. The supernatant was removed and packing efficiency was assessed by UV-Vis absorbance measurements of the supernatant at 280 nm. The rotor, filled with an estimated 13.5 mg of microcrystalline U-$^{13}C$,$^{15}N$-N-FAT10-C0, was closed with a Vespel cap.

**Lyophilized/rehydrated N-FAT10-C0**

After concentration to approximately 8 mg/mL, U-$^{13}C$,$^{15}N$-N-FAT10-C0 was lyophilized, ground, and packed into a 3.2 mm MAS rotor. The rotor containing an estimated 14.5 mg of protein was closed with a Vespel cap. Rehydration was accomplished by adding Milli-Q water in amounts of 1 µL while monitoring the $^{1}H$-$^{13}C$ cross-polarization spectrum. After addition of a total of 7 µL, no further improvement in spectral resolution was observed.

**N-FAT10-C0 in complex with NUB1L**

Complex formation of N-FAT10-C0 and NUB1L was verified with size-exclusion chromatography (Figure S1). To prepare the complex for MAS NMR spectroscopy, co-sedimentation into the rotor was used. 1 mL of U-$^{13}C$,$^{15}N$-N-FAT10-C0 at a concentration of 3 mg/mL was mixed with 1 mL of natural abundance NUB1L with a concentration of 20 mg/mL, leading to a molar ratio of 1:1. The packing tool containing this solution was centrifuged at RCF 210000 g and 4 °C. As the holding capacity of the packing tool is only 1 mL, the run was, over the course of several days, repeatedly interrupted to remove supernatant and add the original mixture in full. Packing efficiency was assessed by UV-Vis absorbance measurements of the supernatant at 280 nm. The 3.2 mm rotor was closed with a Vespel cap.

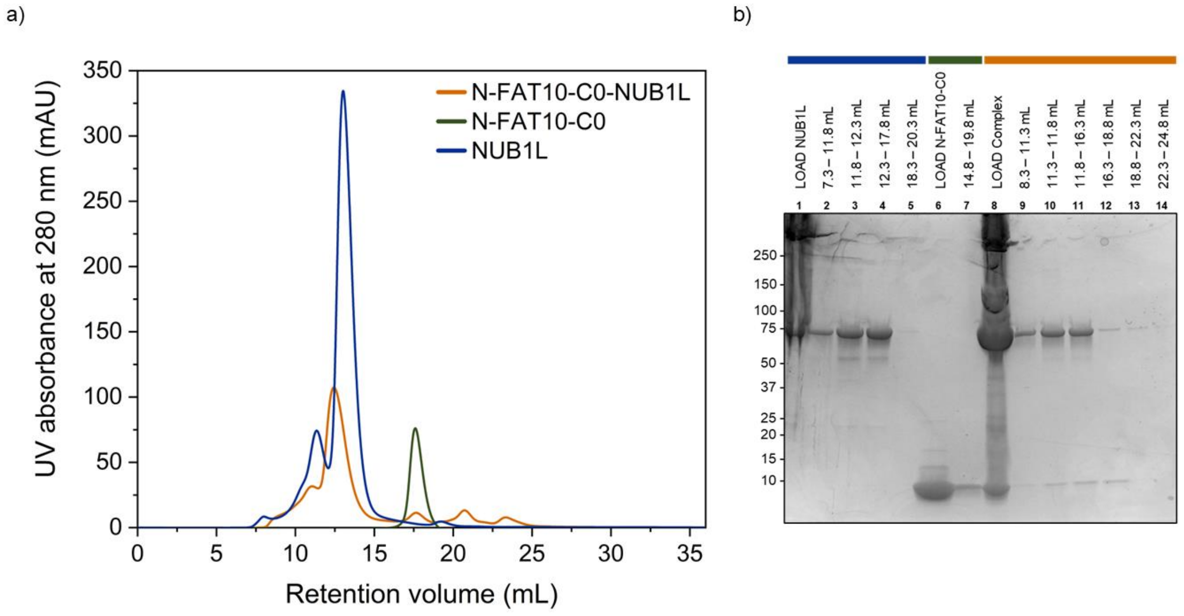

***Figure S1:*** *Size-exclusion chromatography, on a Cytiva ÄKTA pure chromatography system equipped with a Superdex 200 Increase 10/300 GL column, and subsequent SDS-PAGE analysis. (a) Individual runs of isolated N-FAT10-C0 (green) and isolated NUB1L (dark blue) are shown together with a mixture of both N-FAT10-C0 and NUB1L (orange). Molar amounts were adjusted to be the same in each run. The shift to smaller retention volumes in the orange chromatogram indicates a larger molecular weight and thereby complex formation. (b) SDS-PAGE analysis before and after size-exclusion chromatography. 5 µL of gel sample buffer with SDS (reducing conditions) was added to 20 µL of each sample (loads for size-exclusion chromatography as well as collected fractions containing protein). After boiling for 5 min at 95 °C, samples were loaded on a gradient gel and separated in 30 min by applying a constant voltage of 200 V, followed by colloidal Coomassie staining. For molecular weight estimation, Precision Plus Protein Dual Color Standards (Bio-Rad) were used.*

#### 3. MAS NMR spectroscopy

MAS NMR experiments were performed at 18.8 T ($^{1}$H Larmor frequency of 800.3 MHz) in the NMR Core Facility at the University of Konstanz and at 20.0 T ($^{1}$H Larmor frequency of 850.2 MHz) at ETH Zürich. The console was Bruker Avance NEO or Avance III. Bruker 3.2 mm E-free HCN MAS probes were used, together with regular-wall 3.2 mm ZrO rotors (maximum sample volume 32.1 µL). During experiments, the sample was kept at a temperature of approximately 4 °C, unless noted otherwise. Calibration of the Bruker Cooling Unit (BCU) II was performed by monitoring the chemical shift of $^{79}$Br in KBr powder.[3] The carbon dimension was referenced indirectly to DSS in $D_2O$ (0.5 % by weight),[4] i.e., the $^{13}$C adamantane methylene peak is observed at 40.49 ppm. The nitrogen dimension was referenced to liquid ammonia at 25 °C.[5] Table S1 provides an overview of the experiments performed per sample.

***Table S1:*** *MAS NMR spectra obtained for each sample.*

| Experiment | Field (T) | Number of scans | $^{15}$N-$^{13}$C /$^{13}$C-$^{13}$C mixing (ms) | Temperature (°C) | MAS frequency (kHz) | Measurement time |
|---|---|---|---|---|---|---|
| **Microcrystalline U-$^{13}$C,$^{15}$N-N-FAT10-C0** | | | | | | |
| hC | 18.8 | 256 | | 4 | 14.5 | |
| hN | 18.8 | 128 | | 4 | 14.5 | |
| DARR | 18.8 | 32 | 10 | 4 | 14.5 | 16 h |
| ZF TEDOR | 18.8 | 32 | 0.52 | 4 | 14.5 | 7 h |
| ZF TEDOR DARR | 18.8 | 4 | 1.1/40 | 4 | 14.5 | 7 d 21 h |
| **Lyophilized/rehydrated U-$^{13}$C,$^{15}$N-N-FAT10-C0** | | | | | | |
| hC | 18.8 | 256 | | 4 | 14.5 | |
| hN | 18.8 | 512 | | 4 | 19.0 | |
| dREDOR CCP | 20.0 | 1024 | | 4 | 19.0 | |
| dREDOR NCP | 20.0 | 512 | | 4 | 19.0 | |
| DARR | 18.8 | 32 | 10 | 4 | 14.5 | 22 h |
| ZF TEDOR | 18.8 | 128 | 0.84 | 4 | 19.0 | 15 h |
| ZF TEDOR DARR | 18.8 | 8 | 0.84/40 | 4 | 19.0 | 5 d 22 h |
| NCOCX | 20.0 | 32 (2x) | 55.6 (CORD) | 4 | 19.0 | 3 d (2x) |
| **U-$^{13}$C,$^{15}$N-N-FAT10-C0 in complex with natural abundance NUB1L** | | | | | | |
| hC | 20.0 | 128/128/256 | | -10/-2/4 | 19.0 | |
| hN | 20.0 | 2048 | | 4 | 19.0 | |
| C | 20.0 | 128 | | -10/-2/4 | 19.0 | |
| dREDOR CCP | 20.0 | 2048 | | 4 | 19.0 | |
| dREDOR NCP | 20.0 | 2048 | | 4 | 19.0 | |
| DARR | 20.0 | 256 | 10 | 4 | 19.0 | 3 d |
| dREDOR DARR | 20.0 | 1536 | 20 | 4 | 19.0 | 5 d 18 h |
| ZF TEDOR | 20.0 | 256 (2x) | 1.26 | 4 | 19.0 | 1 d 12 h (2x) |
| ZF TEDOR DARR | 20.0 | 8 (3x) | 1.26/50 | 4 | 19.0 | 5 d 5 h (3x) |
| NCOCX | 20.0 | 64 | 55.6 (CORD) | -2 | 19.0 | 6 d |

Typical π/2-pulse lengths were 3.0-3.5 µs for $^{1}$H, 3.0-5.0 µs for $^{13}$C and 6.0-10.5 µs for $^{15}$N. $^{1}$H-$^{13}$C and $^{1}$H-$^{15}$N cross-polarization (CP)[6] are realized using a linearly ramped RF field on the $^{1}$H channel (either from 70 or 90 to 100 % amplitude); the centre of the ramp was set to match the n = +1 Hartmann-Hahn condition.[7,8] Swept-frequency two-pulse phase modulation ($SW_f$-TPPM)[9] or small phase incremental alternation with 64 steps (SPINAL-64)[10] were applied for proton decoupling. The States-TPPI method[11] was used for phase-sensitive detection in the indirect dimensions.

The ZF TEDOR DARR sequence provides NCACX and NCOCX backbone connections within one three-dimensional spectrum, but at the cost of large spectral widths. To keep the number of increments in the indirect dimensions under control, it sufficed for microcrystalline N-FAT10-C0 to truncate the free induction decay along the indirect $^{13}$C dimension and pad it using linear forward prediction. For lyophilized/rehydrated N-FAT10-C0 and the N-FAT10-C0/NUB1L complex, we used aliasing. At a spinning frequency of 19.0 kHz, a convenient strategy is to set the dwell time of the indirect $^{13}$C dimension equal to one rotor period and the carrier frequency to approximately 50 ppm.[12] The carbonyl resonances are now observed at the positions of their spinning sidebands just above the aliphatic region.

Before running a $^{15}$N-$^{13}$C-$^{13}$C ZF TEDOR DARR experiment, a $^{15}$N-$^{13}$C ZF TEDOR experiment served to set the acquisition parameters correctly, including the mixing time and the duration of the REDOR π-pulses. The z-filter delays were set to ~200 μs, matching a multiple of the rotor period. Table S2 gives a complete overview of the acquisition parameters per experiment and sample.

For free induction decays with unnecessary long acquisition times, the number of effective points was adjusted accordingly and zero-filling was applied. 1D spectra were baseline corrected, but no apodization was used. For 2D and 3D spectra, 45°- or 72°-shifted squared-sine bell apodization (corresponding to an SSB value of 4 or 2.5 in TopSpin and an offset of 0.25 or 0.4 in NMRPipe) was used. The resolution of the ZF TEDOR DARR spectrum of microcrystalline N-FAT10-C0 and of the dREDOR DARR spectrum of N-FAT10-C0/NUB1L was improved by forward linear prediction to 1.5 and, respectively, 2.0 times the number of the original data points in the indirect carbon dimension.

***Table S2:** Acquisition parameters for MAS NMR experiments.*

| Experiment | CP contact time (ms) | | Radio frequency field strength (kHz) | | | | | | | | Dwell time (μs) | | | Acquisition time (ms) | | | Carrier position (ppm) | | | Recycle delay (s) |
|---|---|---|---|---|---|---|---|---|---|---|---|---|---|---|---|---|---|---|---|---|
| | $^{1}H$-$^{13}C$/ $^{1}H$-$^{15}N$ CP | $^{15}N$-$^{13}C$ DCP | $^{1}H$-$^{13}C$/$^{1}H$-$^{15}N$ CP | | $^{15}N$-$^{13}C$ DCP | | π/2- and π-pulses | | | Decoupling | | | | | | | | | | |
| | | | $^{1}H^{\#}$ | $^{13}C$/$^{15}N$ | $^{15}N$ | $^{13}C^{\#}$ | $^{1}H$ | $^{13}C$ | $^{15}N$ | $^{1}H$ | $^{13}C$ | $^{13}C$ | $^{15}N$ | $^{13}C$ | $^{13}C$ | $^{15}N$ | $^{13}C$ | $^{13}C$ | $^{15}N$ | |
| **Microcrystalline U-$^{13}C$,$^{15}N$-N-FAT10-C0** | | | | | | | | | | | | | | | | | | | | |
| hC | 0.9 | | 67.9$^{r}$ | 50.0 | | | 83.3 | | | 71.4, $SW_f$-TPPM | 8.5 | | | 17.4 | | | 100 | | | 4.0 |
| hN | 1.0 | | 74.8$^{r}$ | 50.0 | | | 71.4 | | | 71.4, SPINAL-64 | | | 12.0 | | | 12.3 | | | 104 | 4.0 |
| DARR | 1.0 | | 75.9$^{r}$ | 50.0 | | | 83.3 | 50.0 | | 71.4, SPINAL-64 | 11.3 | 23.0 | | 23.1 | 6.9 | | 100 | 100 | | 3.0 |
| ZF TEDOR | 0.9 | | 67.9$^{r}$ | 50.0 | | | 83.3 | 50.0 | 23.8 | 71.4, $SW_f$-TPPM | 12.0 | | 69.0 | 13.2 | | 10.3 | 100 | | 104 | 2.5 |
| ZF TEDOR DARR | 0.9 | | 67.9$^{r}$ | 50.0 | | | 83.3 | 50.0 | 23.8 | 71.4, $SW_f$-TPPM/ SPINAL-64 | 11.3 | 23.0 | 69.0 | 13.6 | 2.9 | 8.8 | 100 | 100 | 104 | 2.5 |
| **Lyophilized/rehydrated U-$^{13}C$,$^{15}N$-N-FAT10-C0** | | | | | | | | | | | | | | | | | | | | |
| hC | 0.9 | | 67.9$^{r}$ | 50.0 | | | 83.3 | | | 71.4, SPINAL-64 | 6.1 | | | 25.0 | | | 100 | | | 3.0 |
| hN | 0.7 | | 64.3$^{r}$ | 41.7 | | | 83.3 | | | 71.4, SPINAL-64 | | | 10.0 | | | 20.5 | | | 104 | 3.0 |
| dREDOR CCP | 4.0 | | 63.0$^{r}$ | 38.7 | | | 100 | 50.0 | 48.1 | 98.3, SPINAL-64 | 5.8 | | | 11.9 | | | 100 | | | 2.0 |
| dREDOR NCP | 4.0 | | 70.6$^{t}$ | 46.5 | | | 100 | 50.0 | 48.1 | 98.3, SPINAL-64 | | | 14.6 | | | 29.9 | | | 122 | 2.0 |
| DARR | 0.9 | | 67.9$^{r}$ | 50.0 | | | 83.3 | 50.0 | | 71.4, SPINAL-64 | 11.3 | 23.0 | | 23.1 | 9.3 | | 100 | 100 | | 3.0 |
| ZF TEDOR | 0.8 | | 72.2$^{r}$ | 50.0 | | | 83.3 | 83.3 | 36.2 | 90.6/83.3, $SW_f$-TPPM | 12.0 | | 52.6 | 24.6 | | 5.3 | 100 | | 87 | 2.0 |
| ZF TEDOR DARR | 0.9 | | 72.2$^{r}$ | 50.0 | | | 83.3 | 83.3 | 36.2 | 90.6/83.3, $SW_f$-TPPM; 83.3, SPINAL-64 | 6.1 | 52.6 | 105.3 | 12.5 | 5.0 | 8.4 | 50 | 50 | 87 | 2.0 |
| NCOCX | 0.7 | 1.3 | 79.5$^{t}$ | 41.7 | 24.0 | 56.2$^{t}$ | 83.3 | 62.5 | 41.7 | 90.0, $SW_f$-TPPM; 100 kHz, CW | 11.0 | 210.5 | 210.5 | 22.5 | 5.1 | 8.4 | 85 | 85$^{†}$ | 122 | 2.0 |
| **U-$^{13}C$,$^{15}N$-N-FAT10-C0 in complex with natural abundance NUB1L** | | | | | | | | | | | | | | | | | | | | |
| hC | 1.0 | | 72.7$^{r}$ | 50.0 | | | 83.3 | | | 83.3, SPINAL-64/ $SW_f$-TPPM* | 5.8 | | | 11.9 | | | 100 | | | 3.0 |
| hN | 0.8 | | 75.6$^{t}$ | 48.3 | | | 100 | | | 98.2, SPINAL-64 | | | 14.6 | | | 29.9 | | | 122 | 2.0 |
| C | | | | | | | | 62.5 | | 83.3, $SW_f$-TPPM | 5.8 | | | 11.9 | | | 100 | | | 4.0 |
| dREDOR CCP | 4.0 | | 69.1$^{r}$ | 40.8 | | | 100 | 50 | 47.2 | 98.2, SPINAL-64 | 5.8 | | | 11.9 | | | 100 | | | 2.0 |
| dREDOR NCP | 4.0 | | 75.6$^{t}$ | 48.3 | | | 100 | 50 | 47.2 | 98.2, SPINAL-64 | | | 14.6 | | | 29.9 | | | 122 | 2.0 |
| DARR | 1.0 | | 72.7$^{r}$ | 50.0 | | | 83.3 | 50.0 | | 83.3, SPINAL-64 | 11.0 | 21.1 | | 11.3 | 5.2 | | 100 | 100 | | 2.0 |
| dREDOR DARR | 4.0 | | 69.1$^{r}$ | 40.8 | | | 100 | 50 | 47.2 | 98.2, SPINAL-64 | 11.0 | 21.1 | | 22.5 | 1.6 | | 100 | 100 | | 2.0 |
| ZF TEDOR | 1.0 | | 72.7$^{r}$ | 50.0 | | | 83.3 | 62.5 | 38.5 | 96.2/83.3, $SW_f$-TPPM | 11.0 | | 52.6 | 11.3 | | 6.5 | 100 | | 104 | 2.0 |
| ZF TEDOR DARR | 1.0 | | 72.7$^{r}$ | 50.0 | | | 83.3 | 62.5 | 38.5 | 96.2/83.3, $SW_f$-TPPM; 83.3, SPINAL-64 | 6.1 | 52.6 | 105.3 | 12.5 | 5.0 | 7.4 | 56 | 56 | 83 | 2.0 |
| NCOCX | 0.8 | 1.0 | 80.5$^{t}$ | 50.0 | 11.0 | 9.4$^{t}$ | 83.3 | 50.0 | 41.7 | 90.0, $SW_f$-TPPM; 90 kHz, CW | 11.0 | 210.5 | 210.5 | 22.5 | 5.1 | 8.4 | 85 | 85$^{†}$ | 122 | 2.0 |

$^{\#}$Corresponds to 100 % of the amplitude. $^{r}$Linear ramp. $^{t}$Tangent. *Decoupling sequence for $^{1}H$-$^{13}C$ CP temperature series. $^{†}$For $^{15}N$-$^{13}C$ SPECIFIC-CP, the CO carrier was centred at 174 ppm.

### 4. $^1H$-$^{15}N$ cross-polarization spectra

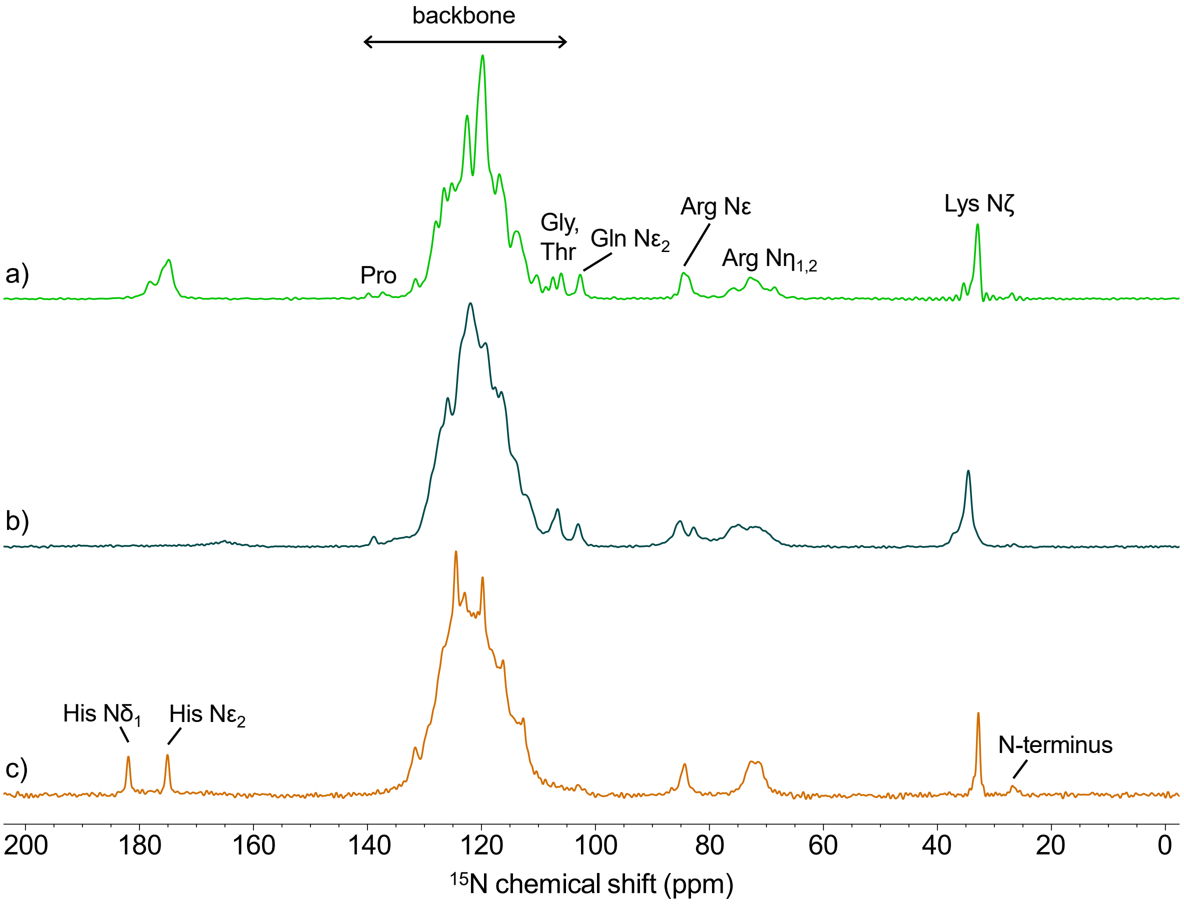

***Figure S2:*** *$^1H$-$^{15}N$ cross-polarization spectra of (a) microcrystalline N-FAT10-C0, (b) lyophilized/rehydrated N-FAT10-C0, and (c) N-FAT10-C0 in complex with NUB1L.*

#### 5. $^{13}C$-$^{13}C$ and $^{15}N$-$^{13}C$ spectra of microcrystalline N-FAT10-C0

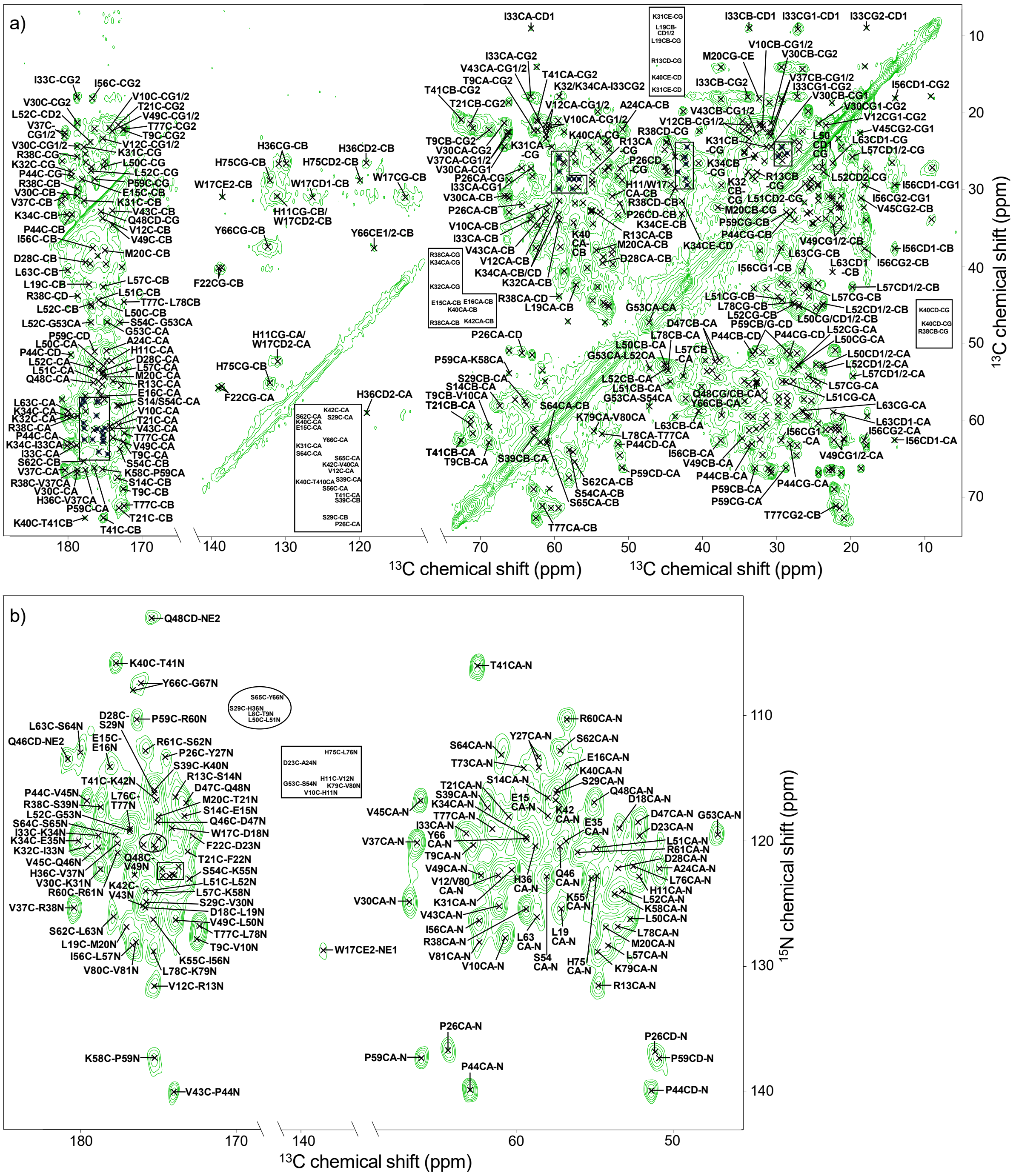

***Figure S3:*** *(a) $^{13}C$-$^{13}C$ DARR spectrum of microcrystalline N-FAT10-C0. The spectrum is dominated by one- and two-bond transfers, but some cross peaks from three-bond transfers are discernible from the noise, e.g. between C' and $C_{\gamma 2}$, $C_{\alpha}$ and $C_{\delta 1}$, and $C_{\gamma 2}$ and $C_{\delta 1}$ of I33 and I53. Some inter-residue contacts are visible, e.g. T9$C_{\beta}$-V10$C_{\alpha}$ and G53$C_{\alpha}$-S54$C_{\alpha}$. To maintain readability, only labels are included of residues that are also observed in the $^{13}C$-$^{13}C$ DARR spectrum of lyophilized/rehydrated N-FAT10-C0, i.e., cross peaks from D18, F22, D23, H36, V45, Q46, K55, K58, R60, R61, T73, H75, L76, K79, V80 and V81 are not labeled in the carbonyl and aliphatic regions. (b) $^{15}N$-$^{13}C$ ZF TEDOR spectrum of microcrystalline N-FAT10-C0.*

### 6. Chemical shifts of microcrystalline N-FAT10-C0

***Table S3:*** *$^{15}$N and $^{13}$C chemical shifts (ppm) of microcrystalline U-$^{13}$C,$^{15}$N-N-FAT10-C0.*

| Residue | N | C' | Cα | Cβ | Cγ | Cδ | Other |
|---|---|---|---|---|---|---|---|
| L8[a] | | 175.3 | 54.9 | 44.3 | 26.1 | 22.2, 21.1 | |
| T9 | 120.3 | 172.6 | 62.8 | 68.8 | 22.2 | | |
| V10 | 127.8 | 174.3 | 60.7 | 31.9 | 22.0, 21.6 | | |
| H11 | 122.8 | 174.3 | 52.2 | 30.8 | 131.1 | 119.0 | |
| V12 | 122.6 | 175.3 | 61.2 | 33.7 | 23.4, 21.6 | | |
| R13 | 131.5 | 173.9 | 54.8 | 32.7 | 27.7 | 43.3 | Nε: 84.7 |
| S14 | 116.5 | 173.4 | 58.0 | 67.4 | | | |
| E15 | 118.0 | 178.1 | 58.0 | 28.6 | 33.4 | | |
| E16 | 114.1 | 175.6 | 56.7 | 28.6 | 33.7 | | |
| W17 | | 174.1 | 52.2 | 30.8 | 114.0 | 126.4, 131.2 | Cε2: 138.6, Cη2: 125.1, Cζ2: 114.4, Nε1: 128.7 |
| D18 | 119.0 | 175.9 | 53.4 | 38.7 | | | |
| L19 | 125.3 | 177.1 | 57.1 | 42.3 | 26.6 | 25.7 | |
| M20 | 126.9 | 175.0 | 54.3 | 37.9 | 32.3 | | Cε: 18.2 |
| T21 | 118.0 | 173.1 | 60.5 | 71.3 | 21.5 | | |
| F22[m] | 120.8 | 172.8 | 55.5 | 40.1 | 139.1 | | |
| D23 | 119.6 | 174.8 | 52.2 | 39.6 | 177.0 | | |
| A24 | 122.3 | 174.7 | 50.9 | 22.1 | | | |
| N25 | 120.7 | 175.0 | 51.2 | 44.3 | | | |
| P26 | 136.7 | 174.6 | 64.4 | 31.8 | 27.2 | 51.2 | |
| Y27[m] | 113.3 | 174.8 | 58.6 | 36.9 | | | Cζ: 156.8 |
| D28[m] | 122.2 | 175.3 | 53.5 | 39.5 | 177.6 | | |
| S29 | 116.2 | 175.8 | 57.5 | 64.0 | | | |
| V30 | 124.8 | 178.7 | 66.8 | 30.8 | 24.4, 21.3 | | |
| K31 | 122.2 | 177.9 | 60.3 | 31.4 | 24.4 | 29.4 | Cε: 42.1, Nζ: 33.0 |
| K32[a] | 120.1 | 179.6 | 59.3 | 33.9 | 27.6 | | |
| I33 | 119.6 | 178.8 | 63.2 | 33.9 | 27.2, 17.9 | 9.1 | |
| K34[a] | 119.5 | 180.1 | 59.4 | 33.1 | 25.9 | 31.4 | Cε: 42.8, Nζ: 35.1 |
| E35 | 120.0 | 175.8 | 56.9 | 28.4 | | | |
| H36[m] | 120.3 | 177.6 | 58.8 | 26.4 | 129.9 | 119.1 | Cε1: 138.2, Nδ1:175.8, Nε2: 174.4 |
| V37 | 120.2 | 180.4 | 66.4 | 30.8 | 23.0, 22.5 | | |
| R38 | 125.4 | 178.7 | 59.4 | 29.8 | 25.7 | 43.8 | Cζ: 159.3, Nε: 83.7, Nη1: 75.8, Nη2: 68.7 |
| S39 | 117.3 | 175.3 | 61.9 | 62.7 | | | |
| K40 | 116.0 | 177.8 | 57.4 | 34.3 | 25.3 | 28.6 | Cε: 42.3, Nζ: 33.1 |
| T41 | 106.0 | 175.2 | 62.4 | 72.6 | 20.9 | | |
| K42 | 116.7 | 176.1 | 57.6 | 29.8 | | | |
| V43 | 125.3 | 174.0 | 61.1 | 32.8 | 22.8 | | |
| P44 | 139.8 | 179.5 | 63.0 | 33.3 | 28.2 | 51.4 | |
| V45 | 116.8 | 178.9 | 66.2 | 31.1 | 22.5, 18.7 | | |
| Q46 | 120.5 | 175.1 | 57.2 | 31.7 | 26.3 | 180.8 | Nε2: 113.5 |
| D47[m] | 118.5 | 173.3 | 52.3 | 38.4 | 175.1 | | |
| Q48 | 117.0 | 176.5 | 55.0 | 32.7 | 32.8 | 175.5 | Nε2: 102.4 |

| | | | | | | | |
|---|---|---|---|---|---|---|---|
| V49 | 122.7 | 173.9 | 62.3 | 34.6 | 22.4, 21.0 | | |
| L50 | 126.2 | 175.2 | 52.8 | 45.0 | 27.0 | 23.9 | |
| L51 | 120.6 | 175.9 | 54.9 | 44.3 | 29.2 | 27.6, 26.2 | |
| L52 | 124.0 | 176.9 | 53.2 | 44.9 | 27.6 | 23.7 | |
| G53 | 119.4 | 174.7 | 47.2 | | | | |
| S54 | 122.7 | 173.0 | 58.1 | 63.7 | | | |
| K55 | 123.0 | 175.4 | 55.3 | 33.8 | 24.4 | 29.3 | Cε: 42.1 |
| I56 | 126.3 | 176.7 | 62.5 | 37.6 | 29.4, 18.2 | 14.1 | |
| L57 | 128.3 | 175.3 | 54.2 | 42.6 | 25.7 | 19.8 | |
| K58 | 124.2 | 175.3 | 53.8 | 32.9 | 25.5 | 28.9 | |
| P59 | 137.3 | 176.4 | 66.1 | 32.9 | 28.7 | 50.9 | |
| R60[m] | 110.3 | 177.7 | 56.8 | 29.7 | 25.3 | | |
| R61 | 120.9 | 175.9 | 56.2 | 31.7 | 28.9 | 43.2 | |
| S62[m] | 112.7 | 177.9 | 57.2 | 65.3 | | | |
| L63 | 126.0 | 180.0 | 58.7 | 40.5 | 26.5 | 22.4, 14.4 | |
| S64 | 113.0 | 177.8 | 61.0 | 62.6 | | | |
| S65 | 119.4 | 175.0 | 61.0 | 62.8 | | | |
| Y66[m] | 119.8 | 176.1 | 59.4 | 37.4 | 132.5 | 132.6 | Cε: 118.1, Cζ: 157.6 |
| G67[m] | 107.4 | | 46.4 | | | | |
| D69[a] | | 173.6 | 58.0 | | | | |
| T73[a] | 114.1 | 172.8 | 59.5 | 71.3 | 21.0 | | |
| H75 | 122.8 | 173.7 | 54.9 | 28.7 | 132.1 | 119.9 | Cε1: 136.6, Nδ1:178.1, Nε2: 174.8 |
| L76 | 122.0 | 176.8 | 52.6 | 45.2 | 27.1 | 24.7 | |
| T77 | 119.0 | 172.4 | 61.6 | 71.0 | 22.0 | | |
| L78 | 126.8 | 175.2 | 53.4 | 44.6 | 27.8 | 23.6 | |
| K79 | 128.8 | 174.0 | 54.8 | 34.4 | 25.1 | | |
| V80 | 122.7 | 176.5 | 61.2 | 33.7 | 22.0 | | |
| V81 | 128.0 | 174.1 | 62.4 | 32.3 | 21.6 | | |

[a]Ambiguous. The backbone $^{15}N$ chemical shifts of K32 and K34 and their succeeding residues are nearly identical, making it difficult to tell them apart. The signals from L8, D69, and T73 are very weak. [m]For these residues, multiple conformations are observed.

### 7. Torsion angle analysis of microcrystalline N-FAT10-C0

**Supporting information for Figure 2d (torsion angle plots)**

- For residues P26-S29, D47, R61, and H75, there is no consensus among the TALOS-N database matches and, hence, no predicted torsion angles are shown.
- No angle predictions could be made for L8, I68-T73, and K82.
- Secondary structure predictions are based on observed chemical shifts except for G67, for which the prediction is based on sequence information.

**Supporting information for Figure 2e (projection of secondary structure per residue)**

- D69 and T73 are ambiguously assigned and colored yellow as secondary structure prediction is based on the sequence.
- Residues K32 and K34 are also ambiguously assigned but colored red (helix) as secondary structure prediction based on (very similar) chemical shifts is highly reliable.
- Residue I74 is unassigned but the secondary structure (light blue = strand) is still predicted based on chemical shifts of neighboring residues.

### 8. $^{13}$C-$^{13}$C and $^{15}$N-$^{13}$C spectra lyophilized/rehydrated N-FAT10-C0

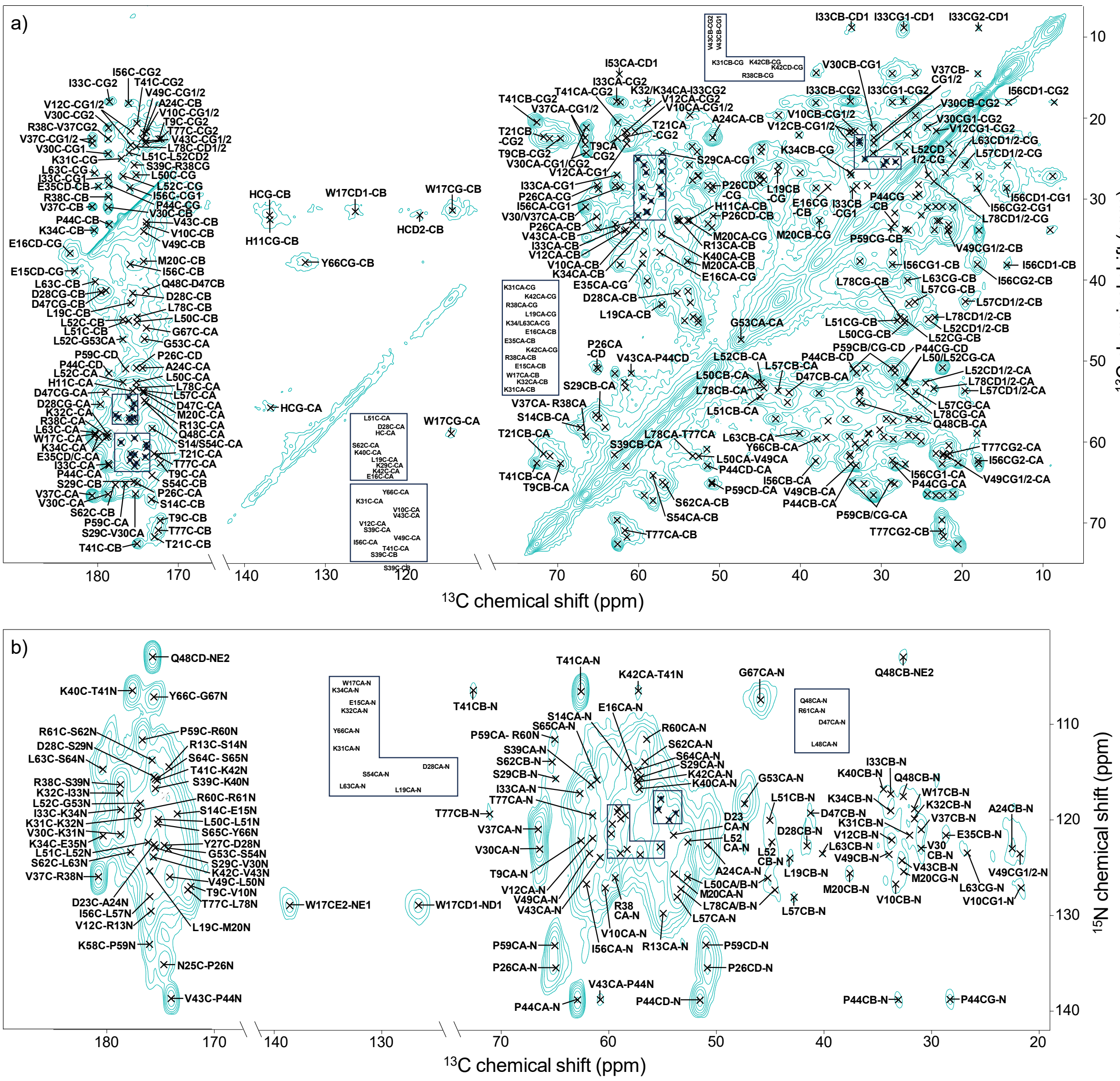

***Figure S4:*** *(a) $^{13}$C-$^{13}$C DARR and (b) $^{15}$N-$^{13}$C ZF TEDOR spectra of lyophilized/rehydrated N-FAT10-C0. Sequentially assigned cross peaks are marked and labeled. As a result of the relatively long mixing time and the large number of scans for the given amount of protein, two- and three-bond N-C$_\beta$ and N-C$_\gamma$ correlations are visible in (b).*

#### 9. Chemical shifts of lyophilized/rehydrated N-FAT10-C0

***Table S4:*** *$^{15}N$ and $^{13}C$ chemical shifts (ppm) of lyophilized/rehydrated U-$^{13}C$,$^{15}N$-N-FAT10-C0.*

| Residue | N | C' | Cα | Cβ | Cγ | Cδ | Other |
|---|---|---|---|---|---|---|---|
| T9 | 122.2 | 172.2 | 62.6 | 69.6 | 22.5 | | |
| V10 | 127.0 | 174.0 | 60.3 | 33.2 | 21.6 | | |
| H11 | 125.4 | 175.7 | 53.7 | 32.6 | 137.0 | | |
| V12 | 121.9 | 175.9 | 61.5 | 33.7 | 23.2, 21.6 | | |
| R13 | 129.7 | 174.2 | 54.9 | 32.7 | | | |
| S14 | 114.3 | 173.4 | 58.2 | 67.2 | | | |
| E15 | 119.5 | 173.4 | 58.5 | 30.2 | 38.8 | 182.9 | |
| E16 | 115.9 | 176.1 | 57.4 | 28.5 | 36.6 | 183.3 | |
| W17 | 119.0 | 180.3 | 59.1 | 31.6 | 114.3 | 126.6, 124.7 | Cε2: 138.5, Nε1: 128.8 |
| L19 | 123.9 | 176.0 | 57.0 | 42.9 | 26.6 | | |
| M20 | 125.4 | 174.4 | 53.9 | 37.7 | 32.6 | | |
| T21 | 118.7 | 173.0 | 61.5 | 71.7 | 22.3 | | |
| D23 | 121.6 | 175.6 | 54.0 | 41.4 | 179.9 | | |
| A24 | 122.7 | 174.5 | 50.8 | 22.4 | | | |
| N25[a] | | 174.6 | | | | | |
| P26[a] | 135.3 | 175.0 | 65.1 | 32.0 | 28.3 | 50.9 | |
| Y27 | | 174.7 | 54.4 | | | | |
| D28 | 122.7 | 175.6 | 55.2 | 41.6 | 179.7 | | |
| S29 | 115.7 | 175.4 | 57.0 | 64.9 | | | |
| V30 | 123.0 | 178.7 | 66.4 | 31.0 | 24.3, 21.1 | | |
| K31 | 121.5 | 177.1 | 60.0 | 32.0 | 25.0 | | |
| K32 | 119.9 | 178.8 | 58.9 | 31.6 | 24.7 | | |
| I33 | 117.3 | 178.7 | 62.7 | 33.8 | 27.1, 18.0 | 8.8 | |
| K34 | 118.9 | 180.3 | 59.2 | 33.9 | 26.6 | 28.4 | |
| E35 | 121.7 | 179.0 | 59.4 | 28.5 | 36.8 | 179.9 | |
| V37 | 120.9 | 180.7 | 66.5 | 30.9 | 23.2, 22.4 | | |
| R38 | 125.9 | 178.7 | 59.3 | 29.7 | 25.7 | | |
| S39 | 116.3 | 175.5 | 61.6 | 62.9 | | | |
| K40 | 116.7 | 177.6 | 57.1 | 34.5 | | | |
| T41 | 106.5 | 175.2 | 62.5 | 72.6 | 20.5 | | |
| K42 | 115.8 | 175.7 | 57.1 | 29.4 | 25.4 | 28.2 | |
| V43 | 124.0 | 174.0 | 60.7 | 32.7 | 23.0, 22.8 | | |
| P44 | 138.9 | 178.6 | 62.9 | 33.1 | 28.3 | 51.5 | |
| D47 | 119.3 | 174.2 | 53.8 | 41.3 | 179.0 | | |
| Q48 | 117.5 | 174.0 | 55.1 | 32.6 | 32.7 | 175.7 | Nε2: 103.0 |
| V49 | 123.5 | 174.2 | 61.8 | 33.9 | 21.8 | | |
| L50 | 126.1 | 175.2 | 52.6 | 45.2 | 27.1 | | |
| L51 | 120.0 | 176.1 | 54.4 | 45.0 | 28.0 | | |
| L52 | 122.6 | 176.9 | 52.7 | 44.9 | 27.2 | 24.2, 24.2 | |
| G53 | 118.6 | 174.3 | 47.3 | | | | |
| S54 | 123.0 | 173.2 | 58.3 | 64.1 | | | |
| I56 | 126.9 | 176.1 | 62.3 | 38.1 | 28.6, 18.1 | 14.5 | |
| L57 | 128.1 | 174.4 | 53.7 | 42.7 | 25.8 | 19.6 | |
| K58[a] | | 176.0 | 52.8 | | | | |

| | | | | | | | |
|---|---|---|---|---|---|---|---|
| P59[a] | 133.2 | 176.6 | 65.0 | 33.5 | 28.8 | 51.0 | |
| R60 | 111.6 | 177.1 | 56.5 | 29.8 | | | |
| R61 | 119.1 | 175.8 | 55.4 | 32.0 | 28.0 | | |
| S62 | 113.9 | 177.8 | 56.6 | 65.3 | | | |
| L63 | 123.5 | 180.4 | 58.9 | 40.1 | 26.7 | 22.1 | |
| S64[a] | 114.7 | 175.5 | 57.3 | 64.7 | | | |
| S65 | 115.9 | 175.3 | 61.1 | 63.3 | | | |
| Y66 | 120.5 | 175.5 | 59.7 | 37.9 | 132.6 | 133.6 | Cε: 118.2, Cζ: 157.5 |
| G67 | 107.1 | 173.8 | 45.9 | | | | |
| T77 | 119.3 | 172.5 | 61.7 | 71.0 | 22.5 | | |
| L78 | 127.3 | 175.2 | 53.3 | 44.6 | 27.7 | 23.5 | |
| K79[a] | 127.3 | | 54.2 | 34.9 | | | |

[a]Ambiguous. Signals from P26 and P59 partially overlap and signals from their preceding residues (N25 and K58) are weak. The signals from S64 overlap with other serines. The signals from K79 are very weak.

### 10. Torsion angle analysis of lyophilized/rehydrated N-FAT10-C0

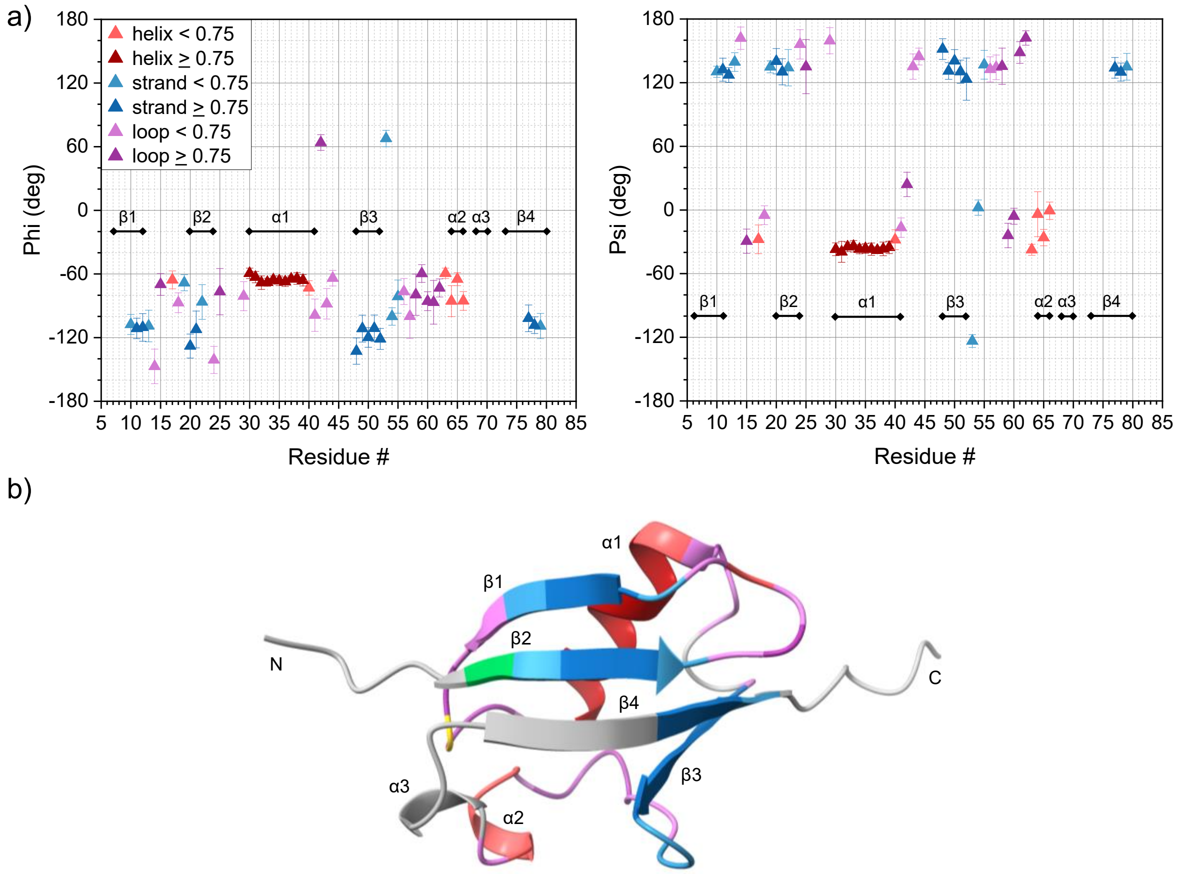

***Figure S5:*** *(a) Backbone torsion angle and secondary structure predictions based on MAS NMR of lyophilized/rehydrated N-FAT10-C0. Error bars correspond to the standard deviation of the Φ and Ψ angles of the best matches in the TALOS-N database. For residues E16, D23, Y27, D28, D47, and G67 there is no consensus among the database matches and, hence, no predicted torsion angles are shown. No angle predictions are made for T9, P26, V45, Q46, I68-L76, and V80. Secondary structure predictions are based on observed chemical shifts and classified as helix, strand, or loop. Predictions with a confidence ≥ 0.75 are highly reliable. (b) Projection of TALOS-N secondary structure predictions per residue on the AlphaFold structure of N-FAT10-C0. D18, F22, H36 and K55 are unassigned, but secondary structure is predicted from the chemical shifts of neighboring residues. For residues N25, K58, P59, S64 and K79, the assignments are ambiguous, but since the secondary structure is reliably predicted from chemical shifts of also neighboring residues, they are color coded accordingly.*

### 11. $^{13}C$-$^{13}C$ and $^{15}N$-$^{13}C$ spectra of N-FAT10-C0 in complex with NUB1L

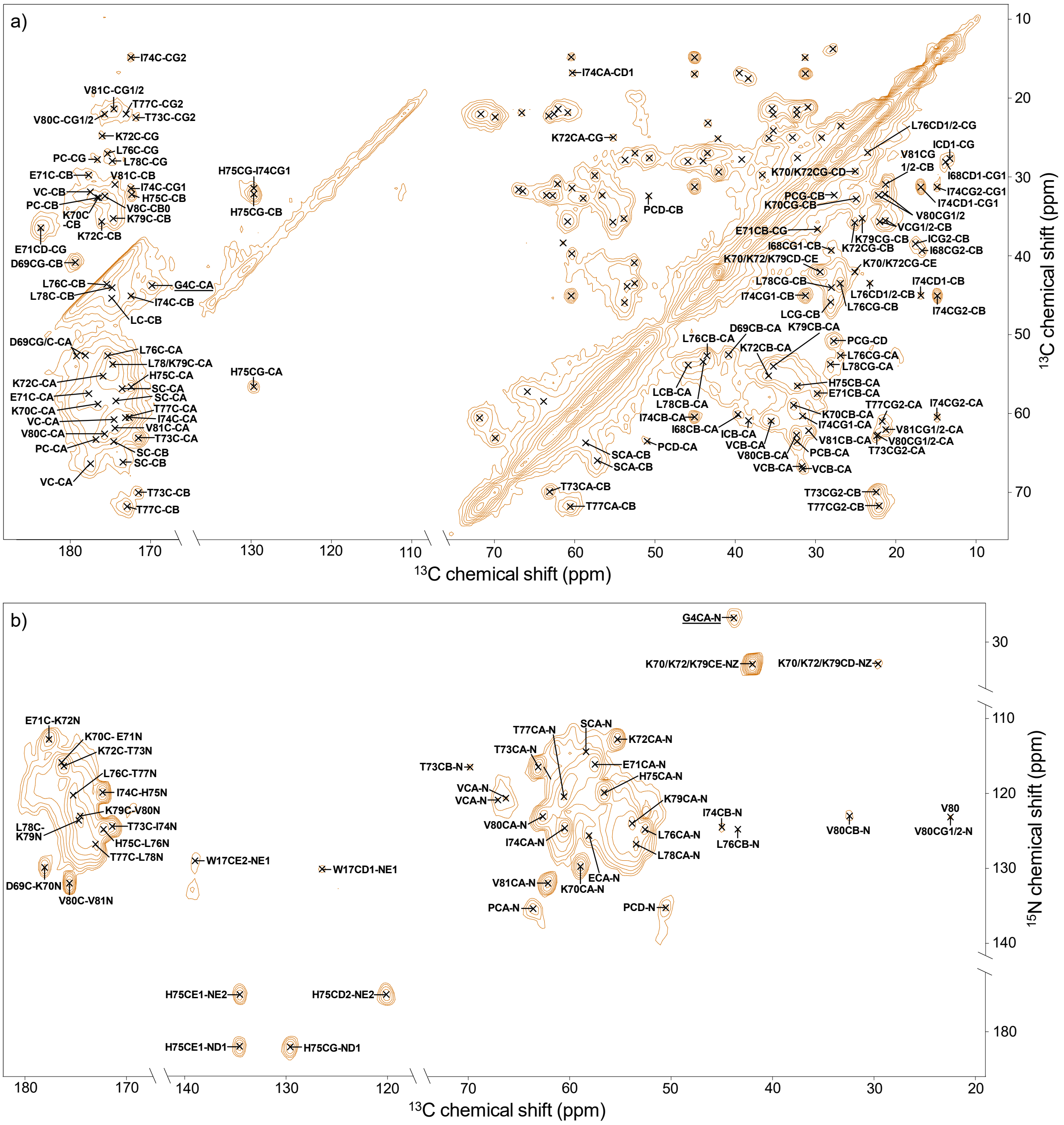

***Figure S6:*** *(a) $^{13}C$-$^{13}C$ DARR and (b) $^{15}N$-$^{13}C$ ZF TEDOR spectra of N-FAT10-C0 co-sedimented with NUB1L. Sequentially assigned cross peaks are marked and labeled.*

#### 12. Temperature dependence of N-FAT10-C0 in complex with NUB1L

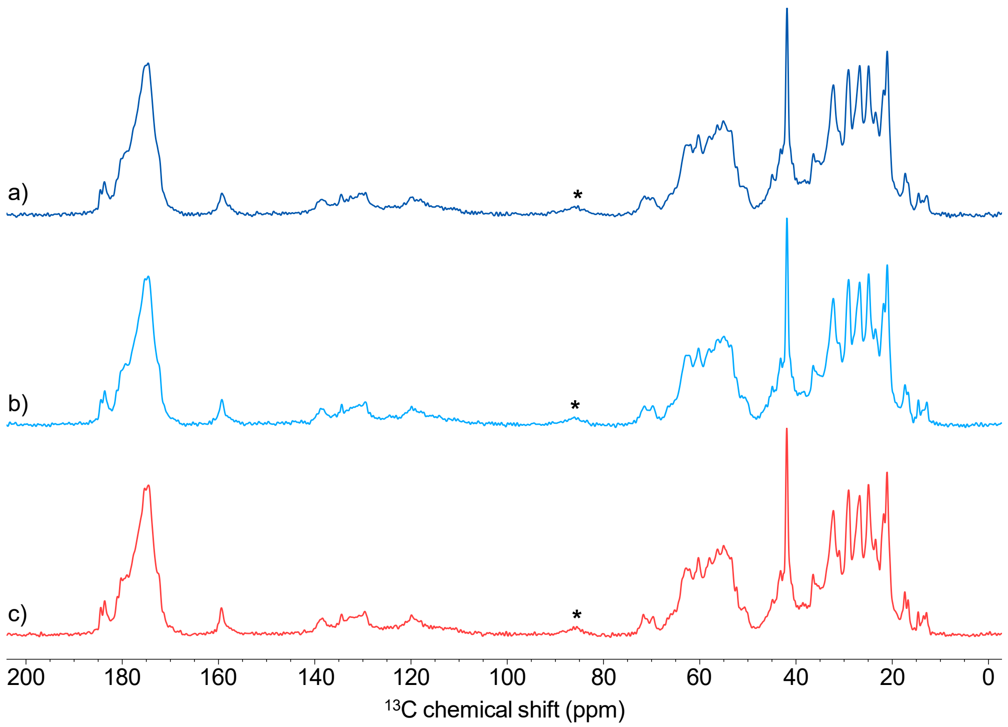

***Figure S7:*** *$^{13}$C direct-excitation spectra of N-FAT10-C0 in complex with NUB1L at temperatures of (a) −10 °C, (b) −2 °C, and (c) 4 °C.*

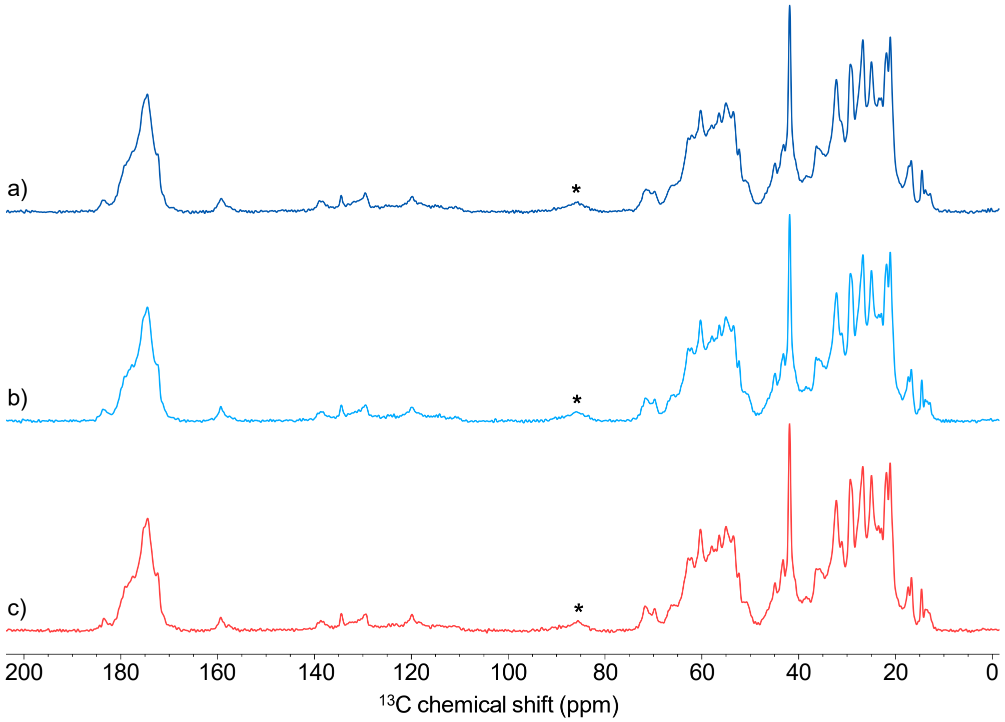

***Figure S8:*** *$^{1}$H-$^{13}$C cross-polarization spectra of N-FAT10-C0 in complex with NUB1L at temperatures of (a) −10 °C, (b) −2 °C, and (c) 4 °C. Compared to the $^{13}$C direct-excitation spectra, the signals in the carbonyl region are relatively weak. The reason is that the cross-polarization contact time of 1 ms does not allow a complete magnetization transfer to $^{13}$C nuclei in this region, which, contrary to most other $^{13}$C nuclei in a protein, do not have a directly bonded $^{1}$H.*

### 13. Chemical shifts of N-FAT10-C0 in complex with NUB1L

***Table S5:*** *$^{15}N$ and $^{13}C$ chemical shifts (ppm) of U-$^{13}C$,$^{15}N$-N-FAT10-C0 co-sedimented with natural abundance NUB1L.*

| Residue | N | C' | Cα | Cβ | Cγ | Cδ | Other |
|---|---|---|---|---|---|---|---|
| G4 | 26.8 | 169.6 | 43.8 | | | | |
| W17 | | | | | | 126.3, 130.7 | Cε2: 138.7, Cη2: 123.8, Cζ2: 114.4, Nε1: 129.6 |
| I68 | | | 60.2 | 39.4 | 27.9, 16.8 | 13.6 | |
| D69 | | 178.1 | 52.6 | 40.8 | 179.2 | | |
| K70 | 129.6 | 176.5 | 58.9 | 32.7 | 25.0 | 29.3 | Cε: 42.0, Nζ: 33.0 |
| E71 | 115.9 | 177.6 | 57.5 | 29.7 | 36.6 | 183.6 | |
| K72 | 112.7 | 176.1 | 55.3 | 35.8 | 25.0 | 29.3 | Cε: 42.0, Nζ: 33.0 |
| T73 | 116.4 | 171.5 | 63.1 | 70.0 | 22.4 | | |
| I74 | 124.4 | 172.5 | 60.4 | 45.1 | 31.3, 14.8 | 16.8 | |
| H75 | 119.7 | 172.4 | 56.5 | 32.2 | 129.6 | 120.2 | Cε1: 134.6, Nδ1: 182.1, Nε2: 175.4 |
| L76 | 124.7 | 175.3 | 52.6 | 43.5 | 26.9 | 23.4, 22.1 | |
| T77 | 120.3 | 172.9 | 60.6 | 71.8 | 22.0 | | |
| L78 | 126.8 | 174.7 | 53.6 | 44.0 | 27.9 | 22.1 | |
| K79 | 123.6 | 174.6 | 53.7 | 35.2 | 24.3 | 29.4 | Cε: 42.0, Nζ: 32.9 |
| V80 | 123.0 | 175.6 | 62.5 | 32.3 | 22.1, 21.3 | | |
| V81 | 131.8 | 174.5 | 62.1 | 31.0 | 21.2 | | |
| E | 125.6 | 181.2 | 58.1 | 31.2 | 36.6 | 184.3 | |
| I | | | 61.2 | 38.5 | 27.6, 17.5 | 13.1 | |
| L | | 174.8 | 53.9 | 45.7 | 27.9 | | |
| P | 135.1 | 176.7 | 63.5 | 32.4 | 27.7 | 50.7 | |
| R | | | | | | 43.4 | Cζ: 159.5, Nε: 85.2, Nη: 71.8 |
| S | 114.4 | 174.5 | 58.3 | 63.8 | | | |
| S | | 173.5 | 57.0 | 66.0 | | | |
| V | 126.5 | 174.3 | 60.9 | 35.6 | 22.0, 21.3 | | |
| V | 120.9 | | 67.0 | 31.7 | | | |
| V | 120.6 | 177.4 | 66.5 | 31.8 | 21.7 | | |
| Y | | | | | | | Cζ: 157.7 |

#### 14. Torsion angle analysis of N-FAT10-C0 in complex with NUB1L

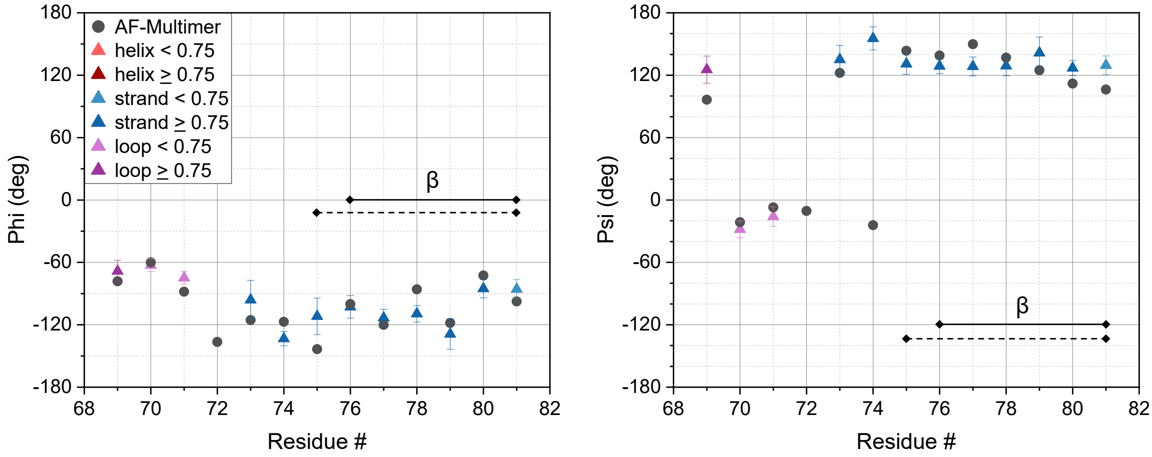

***Figure S9:*** *Backbone torsion angle (triangles) and secondary structure predictions based on MAS NMR of N-FAT10-C0 co-sedimented with NUB1L. For comparison, torsion angles (black dots) of the AlphaFold-Multimer prediction (Figures 5a and S10) are also plotted for comparison. The horizontal solid and dashed black lines indicate the residues classified as* $\beta$*-strand by ChimeraX and VADAR, respectively. Error bars correspond to the standard deviations of the* $\Phi$ *and* $\Psi$ *angles of the best matches in the TALOS-N database. For residue K72, there is no consensus among the database matches and, hence, no predicted torsion angles are shown. All plotted secondary structure predictions are based on observed chemical shifts. Based on sequence information, the secondary structure of I68 is classified as loop. Based on observed chemical shifts, the secondary structure of K72 is classified as loop. Predictions with a confidence* $\geq$ *0.75 are highly reliable.*

#### 15. AlphaFold-Multimer prediction for the interaction of N-FAT10-C0 and NUB1L

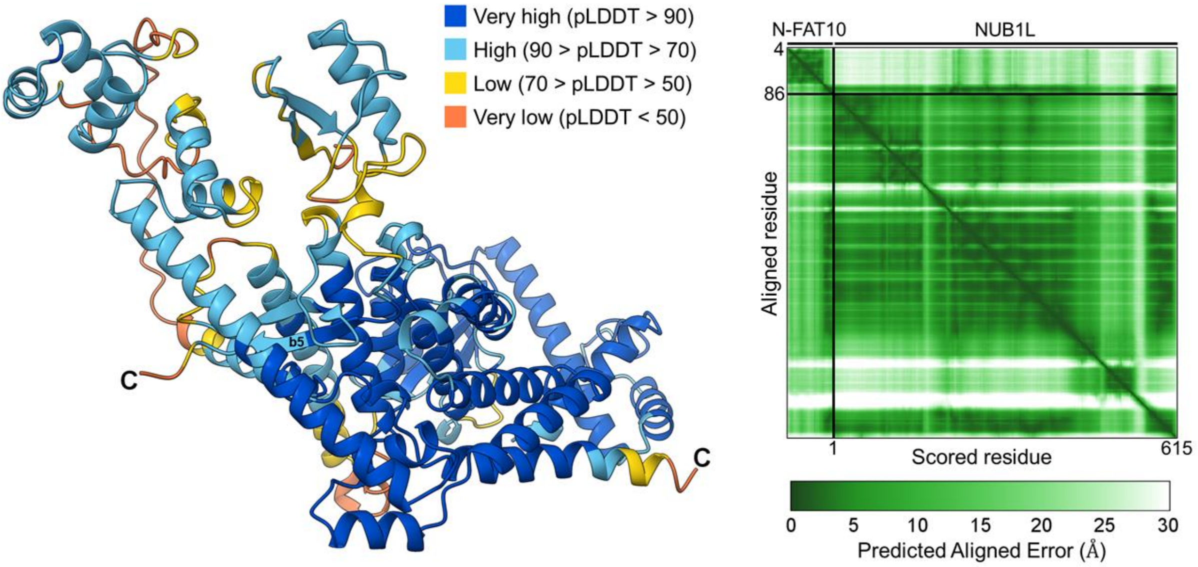

***Figure S10:*** *(left) Best-ranked (0.8·ipTM + 0.2·pTM = 0.78) and relaxed AlphaFold-Multimer structure prediction for the interaction of N-FAT10-C0 and NUB1L. The color code indicates the pLDDT confidence score per residue. The predictions ranked 2-5 were similar, showing the intermolecular, anti-parallel $\beta$-sheet and with model confidence scores above 0.6. All other predictions scored poorly and showed no consistent interaction between N-FAT10-C0 and NUB1L. (right) Plot of the predicted aligned error (PAE). The small positional errors (< 5 Å, dark green) for residues I68 up to S84 of N-FAT10-C0 suggest that NUB1L tightly grasps the $\beta$5-strand.*

### 16. Double-REDOR experiments

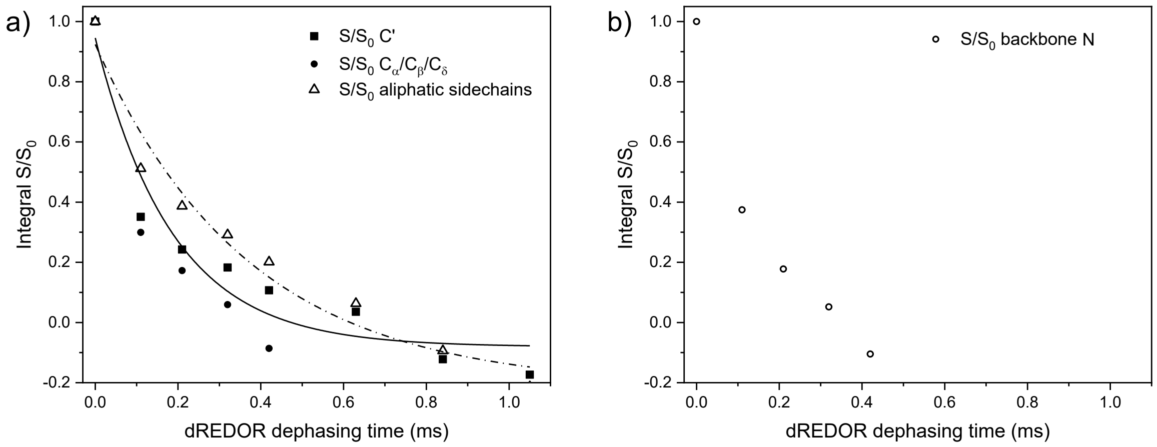

***Figure S11:*** *Double-REDOR dephasing dynamics for (a) different types of* $^{13}C$ *nuclei and (b) backbone* $^{15}N$ *nuclei of isolated lyophilized/rehydrated N-FAT10-C0. Chemical shift ranges over which signal intensity with (*$S$*) and without (*$S_0$*) the double-REDOR dephasing pulses is compared, are indicated in Figures S12 and S13. Exponential decay functions (*$y = y_0 + Ae^{-x/\tau}$*, with* $\tau$ *the double-REDOR dephasing time) fitted to the experimental data points highlight that* $^1H$ *magnetization does not recover after dephasing for the carbonyl and aliphatic sidechain carbons. For the* $C_\alpha$*,* $C_\beta$*,* $C_\delta$ *and the backbone* $^{15}N$ *nuclei, the transverse magnetization of the* $^1H$ *nuclei is lost too quickly after the initial 90°-pulse to properly capture the double-REDOR dephasing curve and it is not possible to fit an exponential decay.*

In their publication on its development, Polenova *et al.* note that the double-REDOR filter struggles to dephase the protons of the $C_\varepsilon$ (42 ppm) of Lys.[13] We observe this behavior not only for this $^{13}C$ nucleus, but also for the $N_\zeta$ (33-34 ppm) of Lys and, in N-FAT10-C0 co-sedimented with NUB1L, the $N_\alpha$ of the N-terminus (26.8 ppm) and the $N_{\varepsilon 2}$ of His (175.4 ppm). We suspect that fast rotation of the ammonium group and/or $^1H$ chemical exchange are at fault. The dephasing behavior of the $N_{\delta 1}$ of His (182.1 ppm) could not be monitored, because the transverse $^1H$ magnetization is very quickly lost.

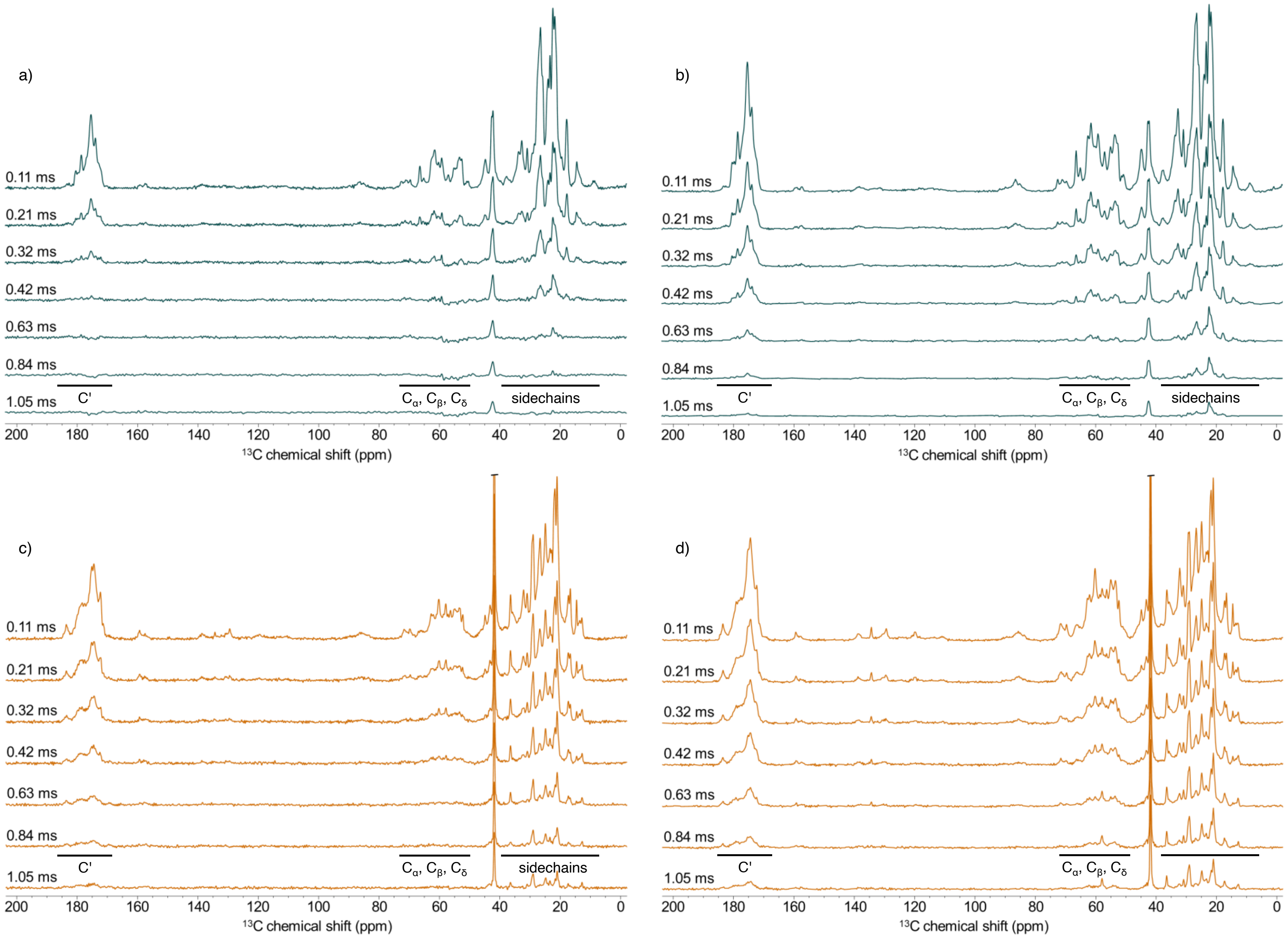

***Figure S12:*** *$^{1}$H-$^{13}$C cross-polarization spectra of lyophilized/rehydrated N-FAT10-C0 (a,b) and of N-FAT10-C0 in complex with NUB1L (c,d) with the double-REDOR dephasing scheme inserted between the initial 90°-pulse and the contact period. The durations of the double-REDOR dephasing are indicated for each spectrum. In the spectra on the right (b,d), the power of the double-REDOR pulses is set to zero.*

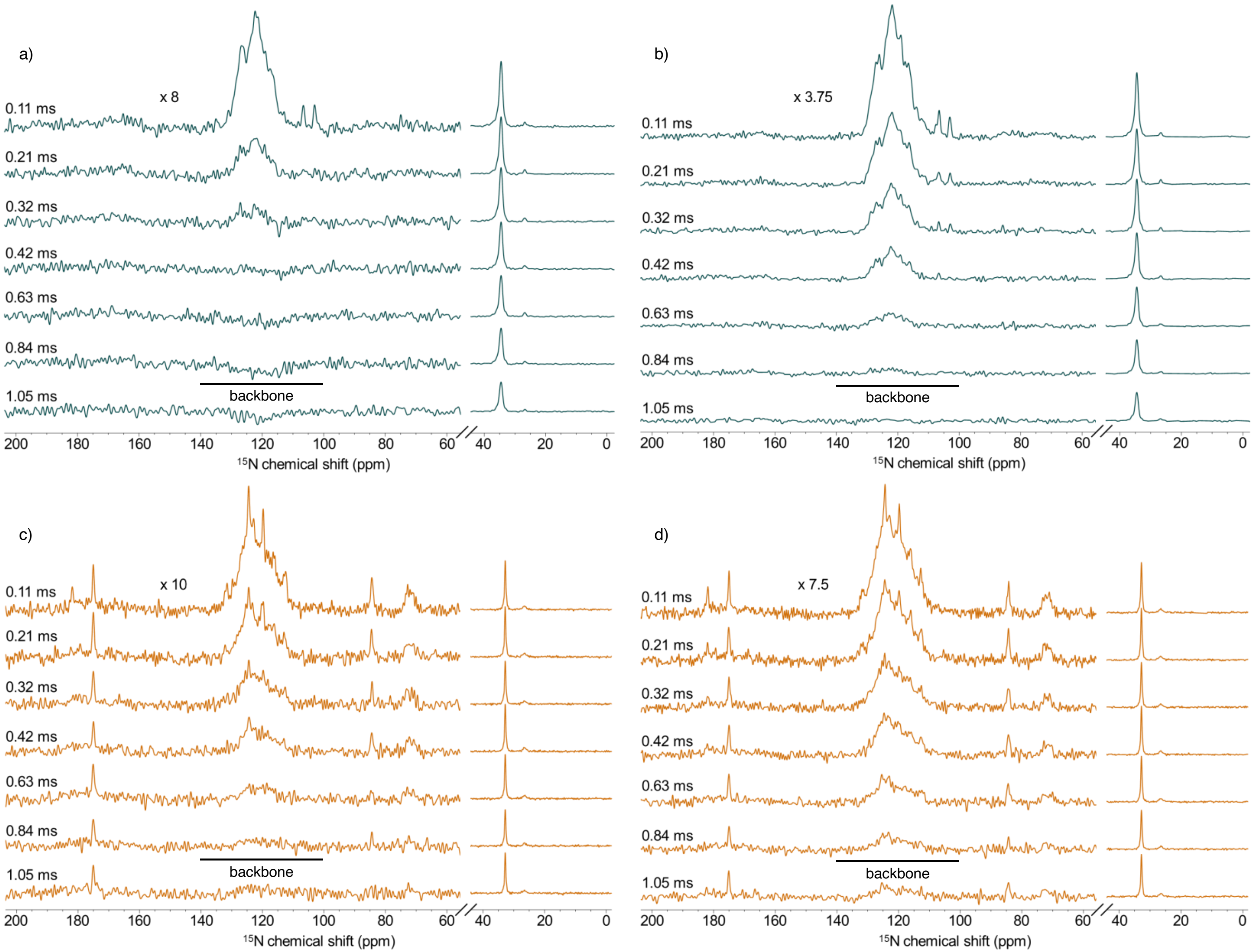

***Figure S13:*** *$^{1}$H-$^{15}$N cross-polarization spectra of lyophilized/rehydrated N-FAT10-C0 (a,b) and of N-FAT10-C0 in complex with NUB1L (c,d) with the double-REDOR dephasing scheme inserted between the initial 90°-pulse and the contact period. The durations of the double-REDOR dephasing are indicated for each spectrum. In the spectra on the right (b,d), the power of the double-REDOR pulses is set to zero.*

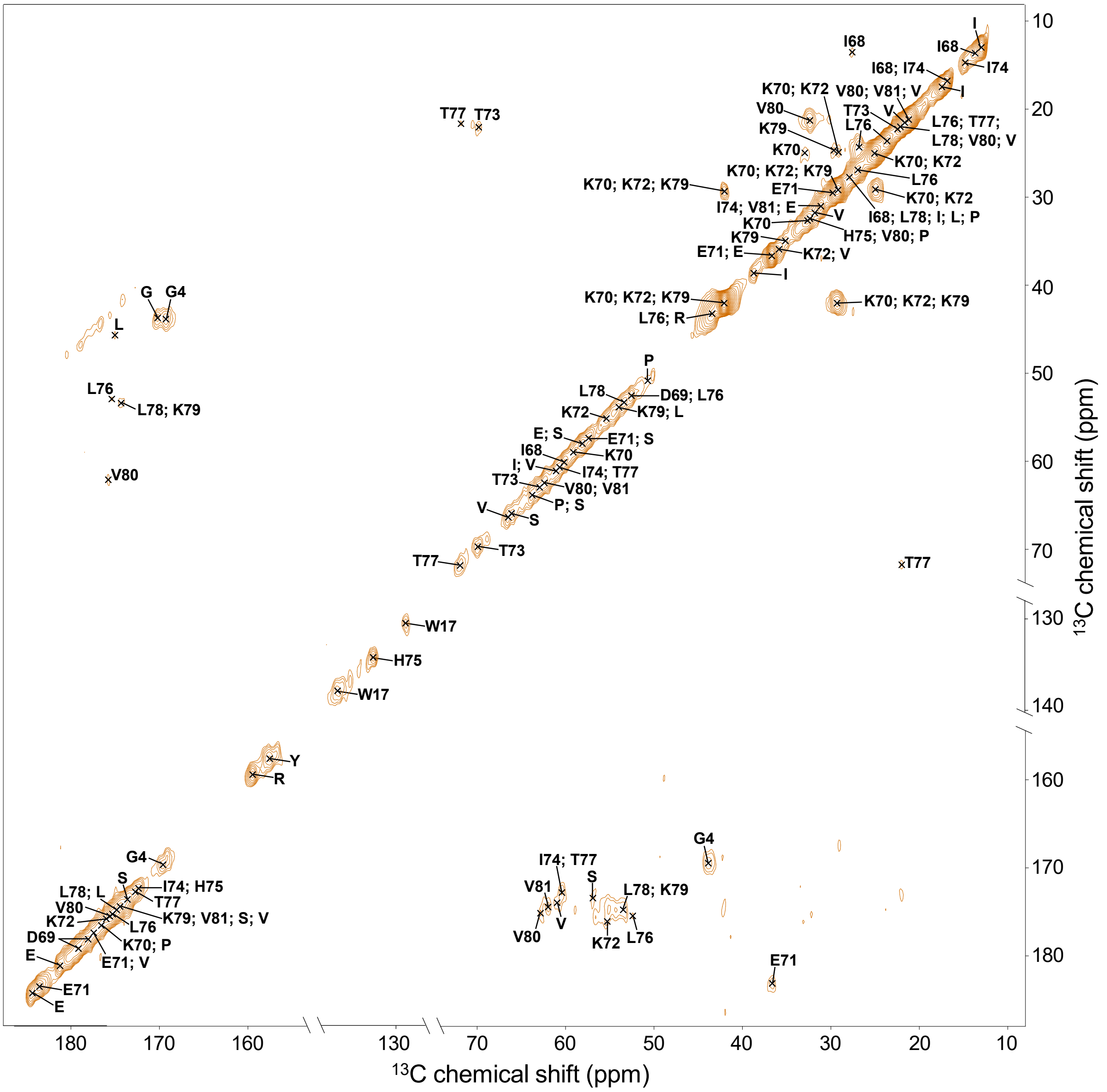

***Figure S14:*** *$^{13}C$-$^{13}C$ DARR spectrum of N-FAT10-C0 co-sedimented with NUB1L recorded after double-REDOR dephasing. All residues that are observed in regular $^{13}C$-$^{13}C$ and $^{15}N$-$^{13}C$ correlation spectra are observed in this spectrum as well, indicating close proximity to the non-dephased $^{1}H$ spins of NUB1L. The asymmetry of the cross-peak patterns across the diagonal is likely due to the weak replenishing of the proton magnetization in the $C_\alpha$-region, i.e., magnetization is transferred from protons via C' to $C_\alpha$ nuclei, but much less via $C_\alpha$ to C' nuclei. The difficulty to dephase the $N_\alpha$ of the N-terminus is likely the reason that the C'-$C_\alpha$ cross peak of G4 at 169.6, 43.8 ppm shows up more strongly (and with a peak doubling of uncertain origin) than the $C_\alpha$-C' cross peak at 43.8, 169.6 ppm on the other side of the diagonal.*
